# Efferocytosis perpetuates substance accumulation inside macrophage populations

**DOI:** 10.1101/583484

**Authors:** Hugh Z. Ford, Lynda Zeboudj, Gareth S. D. Purvis, Annemieke ten Bokum, Alexander E. Zarebski, Joshua A. Bull, Helen M. Byrne, Mary R. Myerscough, David R. Greaves

## Abstract

In both cells and animals, cannibalism can transfer harmful substances from the consumed to the consumer. Macrophages are immune cells that consume their own dead via a process called cannibalistic efferocytosis. Macrophages that contain harmful substances are found at sites of chronic inflammation, yet the role of cannibalism in this context remains unexplored. Here we take mathematical and experimental approaches to study the relationship between cannibalistic efferocytosis and substance accumulation in macrophages. Through mathematical modelling, we deduce that substances which transfer between individuals through cannibalism will concentrate inside the population via a coalescence process. This prediction was confirmed for macrophage populations inside a closed system. We used image analysis of whole slide photomicrographs to measure both latex microbead and neutral lipid accumulation inside murine bone marrow-derived macrophages (10^4^ – 10^5^ cells) following their stimulation into an inflammatory state *ex vivo.* While the total number of phagocytosed beads remained constant, cell death reduced cell numbers and efferocytosis concentrated the beads among the surviving macrophages. Since lipids are also conserved during efferocytosis, these cells accumulated lipid derived from the membranes of dead and consumed macrophages (becoming macrophage foam cells). Consequently, enhanced macrophage cell death increased the rate and extent foam cell formation. Our results demonstrate that cannibalistic efferocytosis perpetuates exogenous (e.g. beads) and endogenous (e.g. lipids) substance accumulation inside macrophage populations. As such, cannibalism has similar detrimental consequences in both cells and animals.

## 1 Introduction

Tissue homeostasis and inflammation resolution require macrophages to phagocytose pathogens and apoptotic cells (efferocytosis), including their own apoptotic macrophages (cannibalistic efferocytosis) [1–3]. Tissues that accumulate harmful stimuli (e.g. pathogens and necrotic cells) become inflamed and populated by large numbers of macrophages and other immune cells. Macrophage numbers increase via the recruitment and differentiation of monocytes from the bloodstream, and the proliferation of tissue-resident or monocyte-derived macrophages; they decrease via apoptosis (primarily) and emigration from the tissue [4–7]. Macrophages regulate inflammation via cytokine signalling and phagocytosis. These processes are primarily mediated by cytoplasmic or cell-surface pattern recognition receptors (PRRs) that detect pathogen- and damage-associated molecular patterns (PAMP/DAMPs) of pathogens and damaged cells [8]. In the presence of PAMP/DAMPs and cytokines, macrophages polarise into a spectrum of pro- and anti-inflammatory states (e.g. M1 and M2) and produce cytokines that orchestrate inflammation amplification and resolution [9–11]. The persistence of PAMP/DAMPs in the tissue or inside macrophages can cause chronic inflammation associated with disease [12–14]. The accumulation of pathogens and sterile substances inside macrophages is a hallmark of a variety of inflammatory diseases [15–19]. For example: *Mycobacterium tuberculosis* infections [20], neutral lipids and cholesterol crystals during atherosclerosis [16, 21, 22], monosodium urate and calcium pyrophosphate dihydrate crystals during gout and pseudogout [17], amyloid-*β* during Alzheimer’s disease [18] and silica and asbestos during inflammation of the lung [19, 23].

Inflammatory macrophages are short lived phagocytes and, as such, gain substances (e.g. pathogens) from the dead macrophages which they consume, i.e., via cannibalistic efferocytosis [20, 24, 25]. Cannibalism, as seen in tumours, can be a beneficial mechanism that allows organisms to scavenge nutrients when the supply is low [26–28]. However, as seen in ecosystems, cannibalism can also transfer harmful substances (e.g. pathogens) between individuals, potentially allowing them to reach toxic levels inside individuals [29]. As such, cannibalism can perpetuate disease transmission in both cell systems (e.g. tuberculosis [20, 24]) and ecosystems (e.g. Kuru neurodegenerative disorder and bovine spongiform encephalopathy [30]). It is conceivable that cannibalism might play a broad pathological role in macrophages as it does across the animal kingdom.

In this study we use experimental and mathematical approaches to elucidate the relationship between cannibalitic efferocytosis and substance accumulation in macrophages. We observe that efferocytosis fuses the contents of two macrophages into one whereas division splits the contents of one macrophage between two. We derive a mathematical model (a coalescence process [31]) based on these observations. We then experimentally verify predictions from the mathematical model. Specifically, our experiments show that efferocytosis coalesces exogenous beads (derived from phagocytosis) and endogenous lipid (derived from the membranes of dead and consumed macrophages) inside murine macrophages following their stimulation into an inflammatory state *ex vivo*. The total quantity of indigestible substances contained inside macrophages is conserved during cannibalistic efferocytosis and thus concentrates inside the population as it decays in size via cell death. Consequently, stimulation of cell death increased the rate and extent of lipid accumulation in macrophages.

## 2 Results

### 2.1 Theoretical considerations

#### 2.1.1 Efferocytosis concentrates substances inside macrophage populations via a coalescence process

To better understand how substances redistribute among macrophages, we tracked the fate of intracellular beads as macrophages underwent apoptosis, efferocytosis and division (see Methods). The time-lapse experiment of Figure 1A shows that beads are conserved during apoptosis and are gained by the consumer cell during efferocytosis. Conversely, Figure 1B shows that beads contained inside one cell are split between daughter cells during cell division.

**Figure 1:**
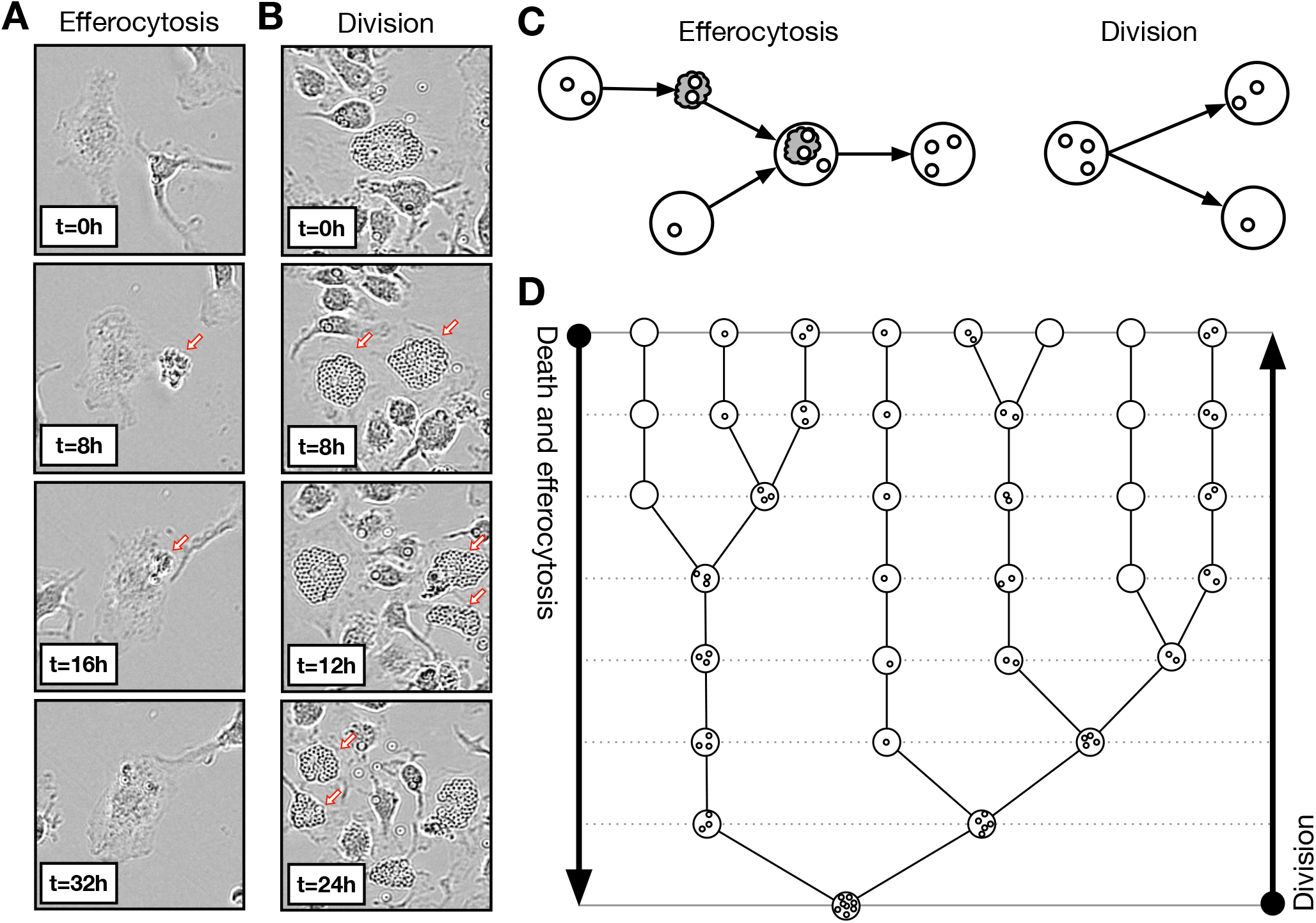
Efferocytosis fuses the contents of two cells into one. **(A)** Time-lapse microscopy of macrophage apoptosis (during times 0 and 8 hours) and efferocytosis (during 16 and 32 hours). The dead macrophage is indicated with an arrow. Intracellular beads are conserved during apoptosis and transferred from the consumed apoptotic cell to live consumer cell during efferocytosis. **(B)** Time-lapse microscopy of macrophage division of one cell (during 0 and 8 hours) and of both daughter cells (during 8 and 12 hours and 12 and 24 hours). The daughter macrophages are indicated with arrows. Intracellular beads are split between daughter cells during division. **(C)** A schematic to illustrate (i) that apoptosis produces one apoptotic cell (grey cloud) from one live cell (white circle) with equal bead content (smaller circles), (ii) efferocytosis transfers the beads content of an apoptotic (consumed) cell to a live (consumer) cell and (iii) division produces two daughter cells whose combined bead content is equal to the bead content of the parent cell. **(D)** A binary tree that represents how apoptosis/efferocytosis decreases the number of cells and increases the number of beads per cell by fusing the bead content of two live cells into one. In reverse, the same binary tree represents how cell division increases the number of cells and decreases the number of beads per cell by splitting the bead content of one cell between two.

These observations cast apoptosis/efferocytosis as a coalescence (or coagulation or aggregation) process that fuses the cargo of two cells into one, and division as a branching (or fragmentation) process that splits the cargo of one cell between two (as illustrated in Figure 1C). As such, bead redistribution within macrophages via apoptosis, efferocytosis and division may be viewed as a coagulation-fragmentation process [32, 33]. Figure 1D depicts a historic lineage of apop-tosis/efferocytosis (binary tree) which illustrates how beads coalesce from several cells with small numbers of beads per cell to one cell with large numbers of beads. When reversed, this binary tree describes how division dilutes bead numbers inside macrophages via a fragmentation process as described previously for other substances [34, 35].

Bead conservation during apoptosis, efferocytosis and division implies that in a closed system the total number of beads within macrophage populations remains constant as the population changes in size. Consequently, the average number of beads per macrophage is inversely proportional to the total number of macrophages, i.e. halving as cell numbers double. More generally, the number of beads per macrophage is coupled to the population dynamics.

#### 2.1.2 A mathematical model of substance accumulation as a coagulation-fragmentation process

To test hypotheses and to facilitate the interpretation of experimental results, we derived and analysed a simple coagulation-fragmentation model for bead accumulation inside macrophages (see Methods and Supplementary Material 1 for details). The model distinguishes live and apoptotic macrophages and keeps track of the number of beads that they contain. We consider macrophage apoptosis, efferocytosis and division (observed *in vitro* and *in vivo)* and neglect monocyte recruitment and cell emigration (observed *in vivo* only). The mass action kinetics used to represent apoptosis, efferocytosis and division are displayed in Figure 2A. Here, we assume conservation of bead numbers. Note that this model describes accumulation of other substances such as lipids, sterile particles or pathogens.

**Figure 2:**
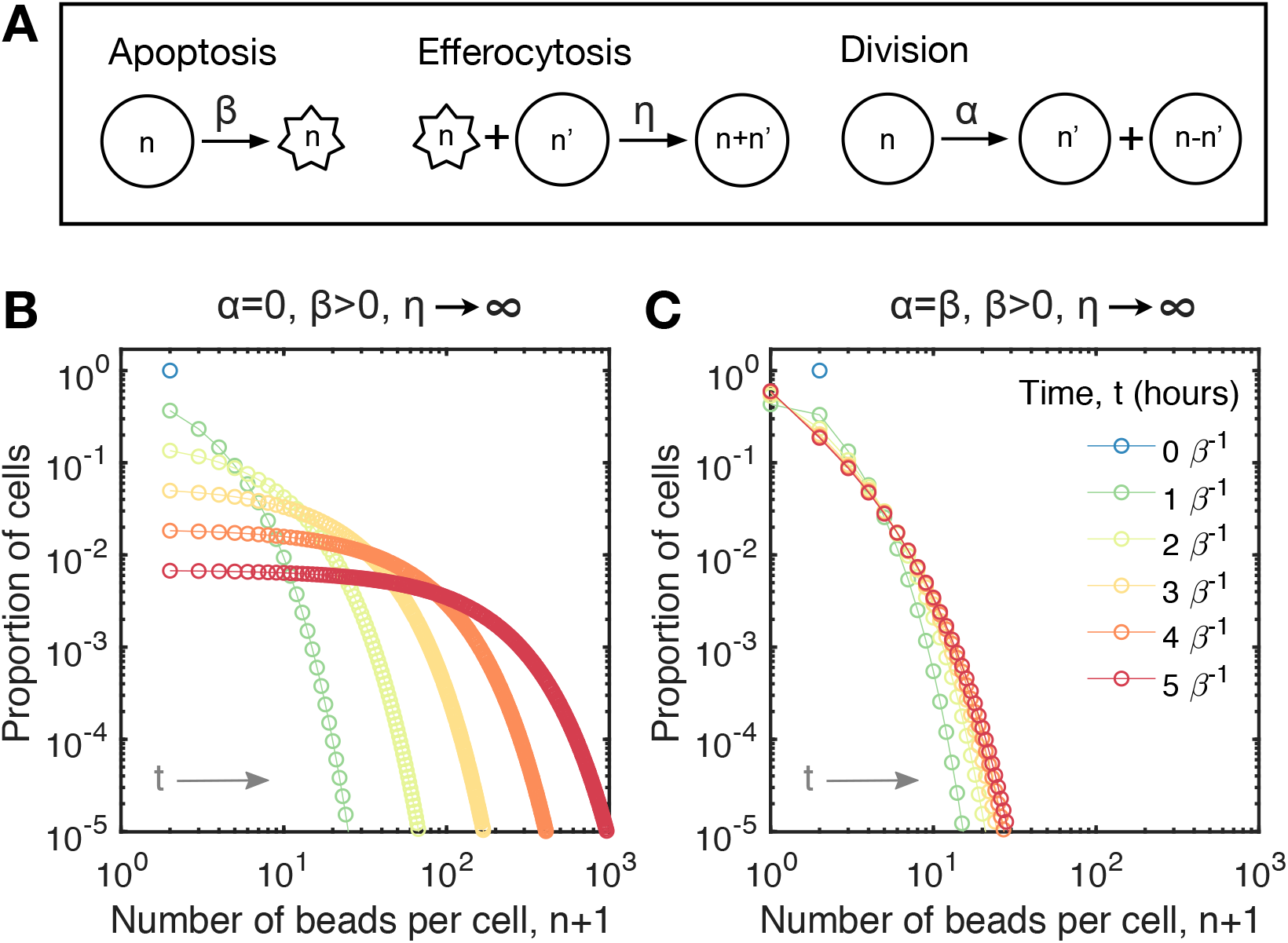
Coagulation-fragmentation model of bead accumulation in macrophages. **(A)** Cartoon schematic of the mass action kinetics for live (circles) and apoptotic (stars) macrophages due to apoptosis, efferocytosis and cell division. We assume: (i) apoptosis occurs at rate *β* (per hour) and converts one live cell with *n* ≥ 0 beads into one apoptotic cell with *n* beads, (ii) efferocytosis occurs at rate *η* (per cell per hour) and transfers *n* beads inside one apoptotic cell (consumed) to one live cell (consumer) with *n*′ ≥ 0 beads to produce one live cell with *n* + *n*′ beads and (iii) division occurs at rate *α* (per hour) and splits *n* beads inside one cell (parent) between two cells (daughters) which contain 0 ≤ *n*′ ≥ *n* beads and *n* – *n*′ beads. **(B)** Model prediction for the time-evolution of the proportion of cells with *n* beads per cell while there is no division (*α* = 0). In this case apoptotic cells are instantly consumed (*η* → ∞) and every cell initially (*N*_0_ cells) contains *n* = 1 bead per cell. The population size *N* declines as *N*(*t*) = *N*_0_*e*^−*βt*^ where *t* represents time (in hours). **(C)** Model prediction for the time-evolution of the proportion of cells with *n* beads per cell while the rates of apoptosis and division balance (*α* = *β*) such that the population size remains constant over time (*N*(*t*) = *N*_0_ cells). In this case apoptotic cells are instantly consumed (*η* → ∞) and every cell initially (*N*_0_ cells) contains *n* = 1 bead per cell. In both (B) and (C), each coloured line represents the predicted distribution after *t* =0, 1, …, 5 macrophage lifespans, *β*^−1^ hours (from blue to red).

Model simulation results for two cases are presented in Figure 2B and C. These cases consider bead accumulation via apoptosis and efferocytosis either with and without cell division. In both cases, we assume that every cell initially contains one bead and apoptotic cells are consumed instantaneously.

Without cell proliferation, the model predicts that beads progressively and heterogeneously accumulate inside the population via efferocytosis as it decays exponentially in size via apoptosis. Figure 2B shows how beads are distributed across the population at different points in time. As cell numbers decrease, this distribution becomes increasingly more uniform across a growing range of bead numbers per cell. This behaviour is characteristic of coalescence processes [36]. Here, the key parameter is the cell death rate. The model predicts that an increased cell death rate enhances bead accumulation via efferocytosis.

If cells initially contain one bead and one unit quantity of endogenous substances (i.e. neutral lipid inside cell membranes), then the number of beads per cell at later time points might also represent the fold increase in the quantity of endogenous substances (which are not degraded or removed) which that macrophage accumulates via efferocytosis. For example, a macrophage with *n* beads might also represent a macrophage that accumulates lipid derived from *n* – 1 other macrophages which have died and then have been consumed.

The model predicts that if cell division and apoptosis rates balance then cell numbers remain constant and the population evolves to an equilibrium where the number of beads per cell remains small. Figure 2C shows how beads are distributed across the population at different points in time. The population tends to an equilibrium because division and apoptosis balance bead dilution/fragmentation and concentration/coagulation effects [37].

### 2.2 Experimental results

#### 2.2.1 Bead accumulation inside macrophages

We designed an experiment to test the prediction of our mathematical model that efferocytosis causes intracellular substances to concentrate inside macrophages. We quantified latex bead (3μm diameter) accumulation within populations of murine bone marrow-derived macrophages (BMDMs) following their *ex vivo* activation into an inflammatory state. Differentiated BMDMs were incubated at a ratio of one latex bead (half red, half blue) per cell and then stimulated with interferon *γ* (IFN*γ*) and lipopolysaccharide (LPS). Unstimulated bead-loaded BMDMs were used as a control. Macrophages stimulated with IFN*γ* and LPS polarise to a classically activated (M1) state found early in acute inflammation and throughout chronic inflammation [11]. Bead accumulation was quantified prior to stimulation then at 24 and 48 hours after stimulation. The number of beads per cell for every cell (10^5^ cells and 10^5^ beads initially) was counted using a new in-house computer image recognition algorithm applied to whole slide photomicrographs (see Methods and Supplementary Material 2). This method was used because M1 macrophages strongly adhere to surfaces which precludes cell resuspention and flow cytometry.

The experiments presented in Figure 3 show that efferocytosis concentrates beads inside macrophages following bead phagocytosis. Figure 3A shows images of bead-loaded BMDMs prior to stimulation (at time 0) and 24 and 48 hours after stimulation with LPS and IFN*γ*. These images illustrate how both the magnitude of, and variation in, the number beads per cell increases as the number of BMDMs decreases. Also shown is an image of unstimulated BMDMs at 48 hours. Unstimulated BMDMs proliferate to high confluency with low numbers of beads per cell. Figure 3B shows that the total number of internalised beads remained constant as the cell population size decreases following stimulation. This is consistent with observation from the pilot experiments shown in Figure 1. We estimate the average death rate of bead-loaded BMDMs stimulated with LPS and IFN*γ* to be 1/60 hours (see Supplementary Material 2). Figure 3C shows that the average number of beads per cell approximately doubled after each day from the initial state of roughly 1 bead per cell. Figure 3D shows how the beads were distributed across the BMDM population 0, 24 and 48 hours after LPS and IFN*γ* stimulation.

**Figure 3:**
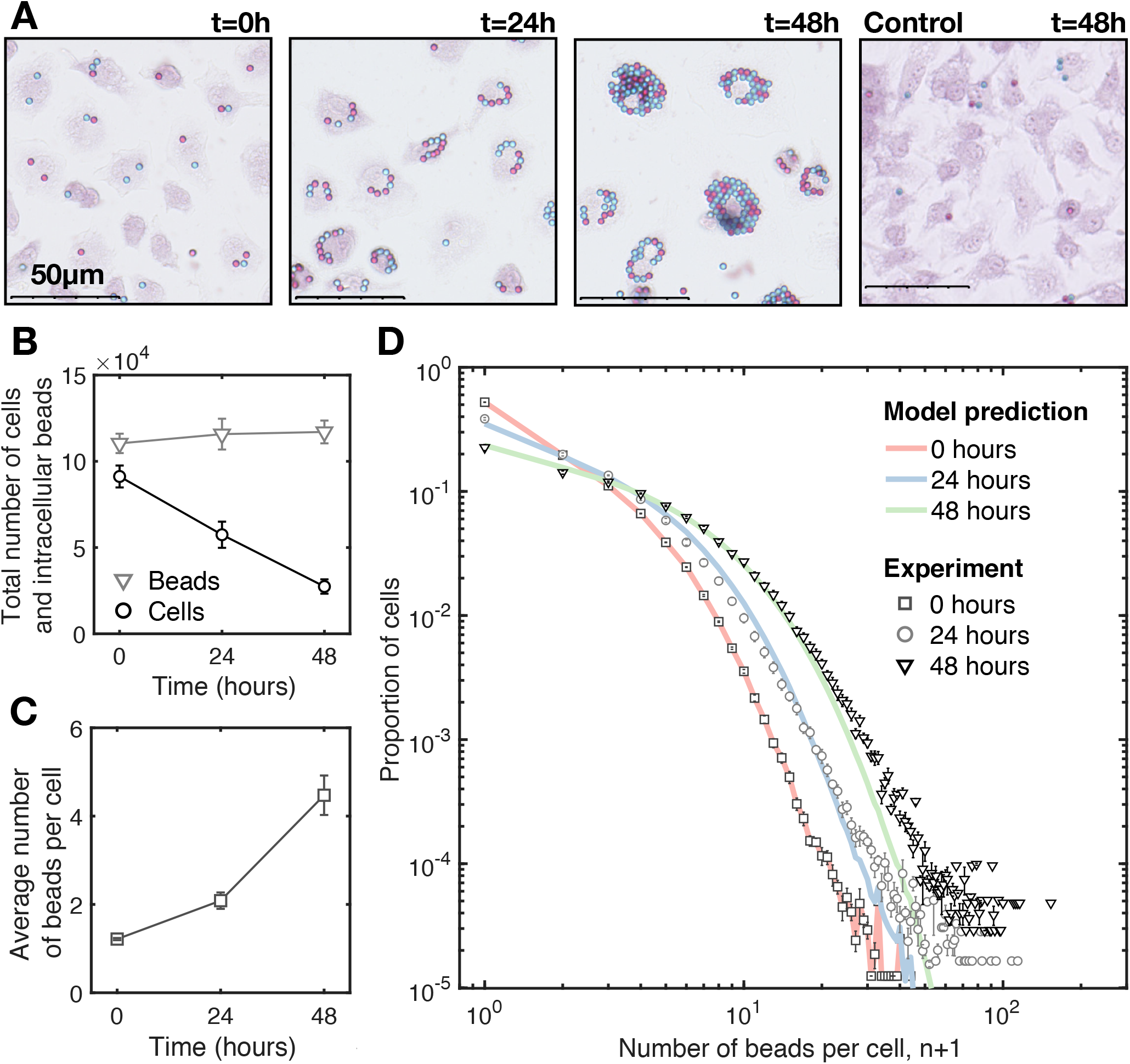
Bead accumulation in M1 macrophages. Quantification of red and blue beads inside safranin-stained murine bone marrow-derived macrophages (BMDMs) following stimulation with IFN-*γ* and LPS. The data presented is averaged from four experiments carried with BMDMs derived from different mice on different days. Initially there were approximately 9 × 10^4^ cells that collectively contain 11 × 10^4^ beads. **(A)** Representative images of BMDMs at 0, 24 and 48 hours after stimulation. Also shown is a representative image of unstimulated BMDMs at 48hours (far right). **(B)** The total number of BMDMs (black circles) and the total number of intracellular beads (grey triangles) over time. **(C)** The average number of beads per cell over time. **(D)** The proportion of BMDMs with number *n* beads per cell prior to stimulation (dark grey squares) and 24 (light grey circles) and 48 (black triangles) hours after stimulation. Also shown are predictions from the mathematical model (no cell division) run from the experimental initial condition (red line and black squares) with death rate 1/60 per hour to 24 (blue line) and 48 (green line) hours; the distribution have r-squared values *R*^2^ = 0.9929 and *R*^2^ = 0.9970 with the experimental distributions at 24 at 48 hours respectively.

The experimental data is consistent with output from the mathematical model (see Methods) when (i) the initial condition of the model, i.e., the distribution of beads across the population, was equal to the experimental data, (ii) bead-loaded BMDMS stimulated with LPS and IFN*γ* die with an average rate of 1/60 per hour, (iii) there is no cell division and (iv) apoptotic cells are consumed instantaneously. Using the method of least squares, we find that mathematical model prediction fits the average values of the experimental data at 24 hours with r-squared value *R*^2^ = 0.9929 and at 48 hours with r-squared value *R*^2^ = 0.9970.

#### 2.2.2 Lipid accumulation inside M1 macrophages

The accumulation of lipid within macrophages (i.e. the generation of foam cells) is a feature of acute and chronic inflammatory responses [22], most notably in atherosclerotic plaques and tuburculosis granulomas [38, 39]. Our mathematical model predicts that efferocytosis causes inflammatory macrophages to accumulate neutral lipid derived from the membranes of dead macrophages that have been consumed by other macrophages. To test this hypothesis experimentally, we use our image recognition algorithm to quantify oil red O-stained lipid droplets inside murine BMDMs stimulated with LPS and IFN-*γ ex vivo.* Unstimulated BMDMs were used as a control. Furthermore, we stimulated BMDMs with LPS and staurosporin (STPN), a compound that promotes apoptosis [40].

The experiments presented in Figure 4 show that neutral lipids accumulate inside BMDMs in the same way that beads accumulate (Figure 2). The images presented in Figure 4A show how the quantity of accumulated lipid per cell increases as the number of cells decreases. For times up until 60 hours, we observe that most apoptotic BMDMs, some of which contain accumulated lipid, are in the process of efferocytosis; most cells were dead after 60 hours. Also shown is an image of unstimulated BMDMs and BMDMs stimulated with LPS and STPN at 48 hours. Unstimulated BMDMs proliferate to high confluency without lipid accumulation. BMDMs stimulated with LPS and STPN accumulate more lipid than BMDMs stimulated with LPS and IFN*γ*. Figure 4B shows that the number of BMDMs declines from approximately 10^5^ (0 hours) to 2 × 10^4^ (60 hours) after LPS and IFN*γ* stimulation, at which time almost all BMDMs stain positive for oil red O. Figure 4C shows that the average lipid content per cell grows exponentially with time, similarly to bead accumulation shown in Figure 3C and consistent with the predictions of our model. If neutral lipid accumulation were due to lipid synthesis or uptake via LDL, then we would expect that the average lipid content per cell would either constantly increase or saturate with time. Figure 4D shows how the the distribution of lipid across the population evolves with time. The extent of lipid accumulation per cell increases over time, i.e., we see cells with larger amounts of lipid. And the cell-to-cell variation with respect to lipid content increases with time, i.e., we see larger disparities in lipid content between cells.

**Figure 4:**
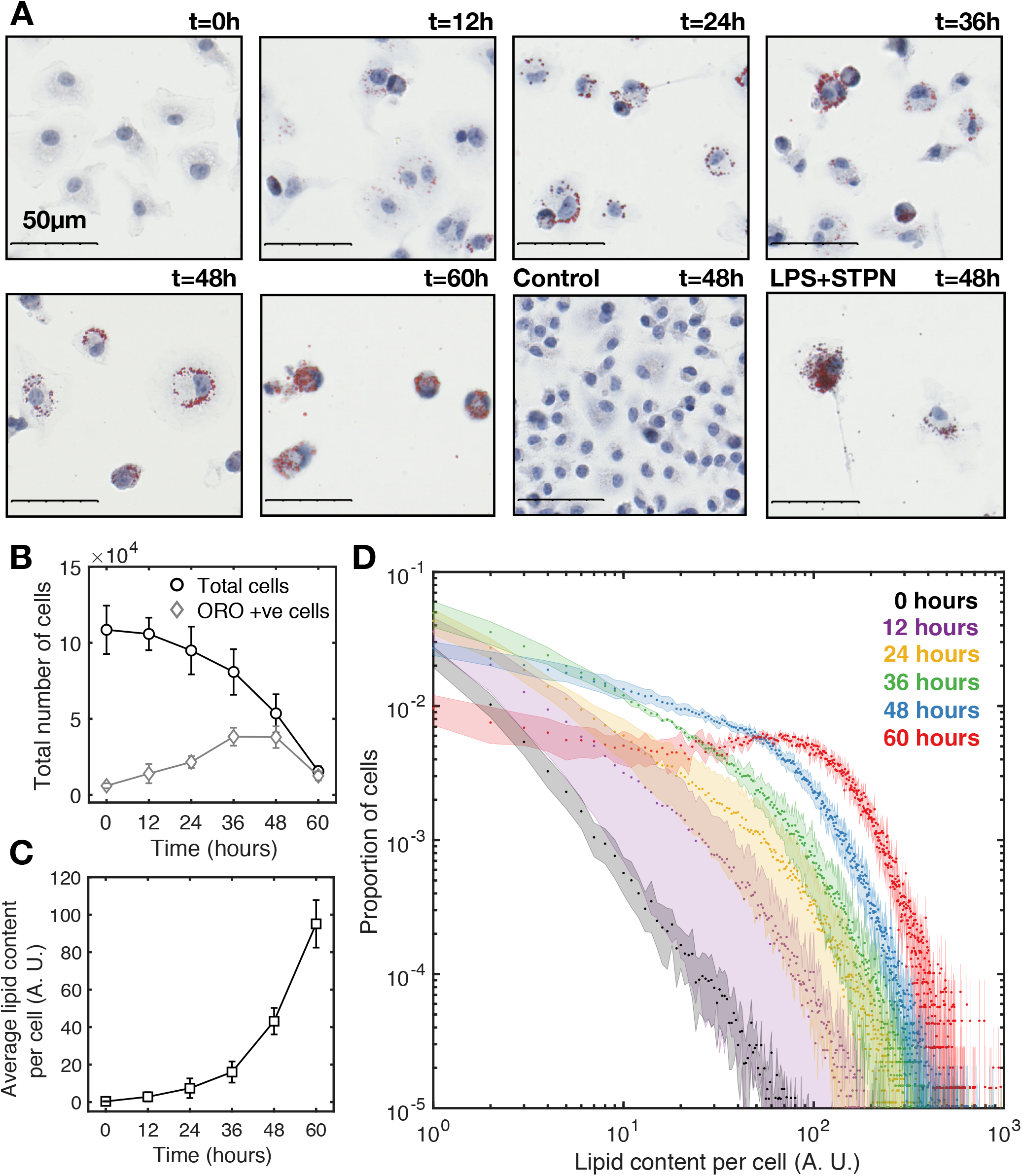
Neutral lipid accumulation in M1 macrophages. Quantification of oil red O-stained neutral lipids inside hematoxylin-stained murine bone marrow-derived macrophages (BMDMs) following stimulation with LPS and IFN-*γ*. Initially there were approximately 1.1 × 10^5^ BMDMs with insignificant quantities of accumulated lipid. The data presented is averaged from four experiments carried with BMDMs derived from different mice on different days. (Continued on the following page.) **(A)** Representative images of BMDMs at 0, 12, 24, 36, 48 and 60 hours after stimulation. Also shown are images of unstimulated BMDMs at 48 hours and BMDMs 48 hours after stimulation with LPS and staurosporin (STPN). **(B)** The total number of BMDMs(black circles) and the number of BMDMs that stain positively for oil red O (grey triangles) over time. **(C)** The average lipid content per cell (arbitrary units) over time. **(D)** The proportion of BMDMs with a range of lipid contents (arbitrary units) at 0 (black), 12 (purple), 24 (yellow), 36 (green), 48 (blue) and 60 (red) hours after stimulation with LPS and IFN-*γ*.

In Figure 4, the quantity of intracellular lipid is expressed in arbitrary units (the cell area that stains positively for oil red O per cell). To facilitate model comparison, these arbitrary units should be expressed in terms of the average endogenous lipid content per macrophage (although this is likely to vary from cell-to-cell).

Like beads, neutral lipids are not degraded during apoptosis and efferocytosis and are unlikely to be degraded by, or removed from, M1 macrophages [22, 41]. As such, the total quantity of neutral lipids (either as components of cellular membranes or lipid droplets) inside the macrophage population should remain constant over time. Efferocytosis essentially translocates the lipid inside the cell membranes (which are not stained by oil red O) of one cell into lipid droplets (which are stained by oil red O) of another cell, as illustrated in Figure 5A. Consequently, the average quantity of accumulated lipid per cell increases as the population size decreases (Figure 4C). This conservation property enables us to scale the arbitrary units of lipids in terms of the average quantity of lipid contained inside macrophage membranes.

Figure 5B shows that in BMDMs stimulated with either LPS and IFN*γ* and LPS and STPN, the amount of accumulated lipid (inside lipid droplets) inside the whole population increases as endogenous lipid (inside cell membranes) decreases such that the total amount of lipid (endogenous+accumulated) remains constant over time. That is, the rate at which endogenous lipid is removed from the population via cell death is equal to the rate at which accumulated lipid is added to the population via efferocytosis. Furthermore, BMDMs stimulated with LPS and STPN die at a faster rate than those stimulated with LPS and IFN*γ*. We estimate the average death rate of BMDMs stimulated with LPS and IFN*γ* to be 1/80 hours and with LPS and STPN to be 1/40 hours (see Supplementary Material 2). Furthermore, BMDMs stimulated with LPS and STPN decay to approximately 25% of the original population size at 48 hours and BMDMs stimulated with LPS and IFN*γ* decay to approximately 50% of the original population size at 48 hours. Consequently, BMDMs stimulated with LPS and STPN display a greater extent of lipid accumulation per cell than those stimulated with LPS and IFN*γ*. This is a prediction from the mathematical model.

Figure 5C shows how lipid is distributed across the BMDM population at 48 hours after stimulation with LPS and IFN*γ* or with LPS and STPN. A larger proportion of BMDMs accumulate excessive quantities of lipid when apoptosis is enhanced. Both distributions are in good qualitative agreement with predictions from the mathematical model. In this modelling case, we assume (i) all cells initially contain one unit of endogenous lipid each (inside cell membranes), (ii) BMDMs stimulated with LPS and IFN*γ* die with an average rate of 1/80 per hour, (iii) BMDMs stimulated with LPS and STPN die with an average rate of 1/40 per hour, (iv) cells do not divide and (v) all apoptotic cells are consumed instantaneously (see Figure 2D).

**Figure 5:**
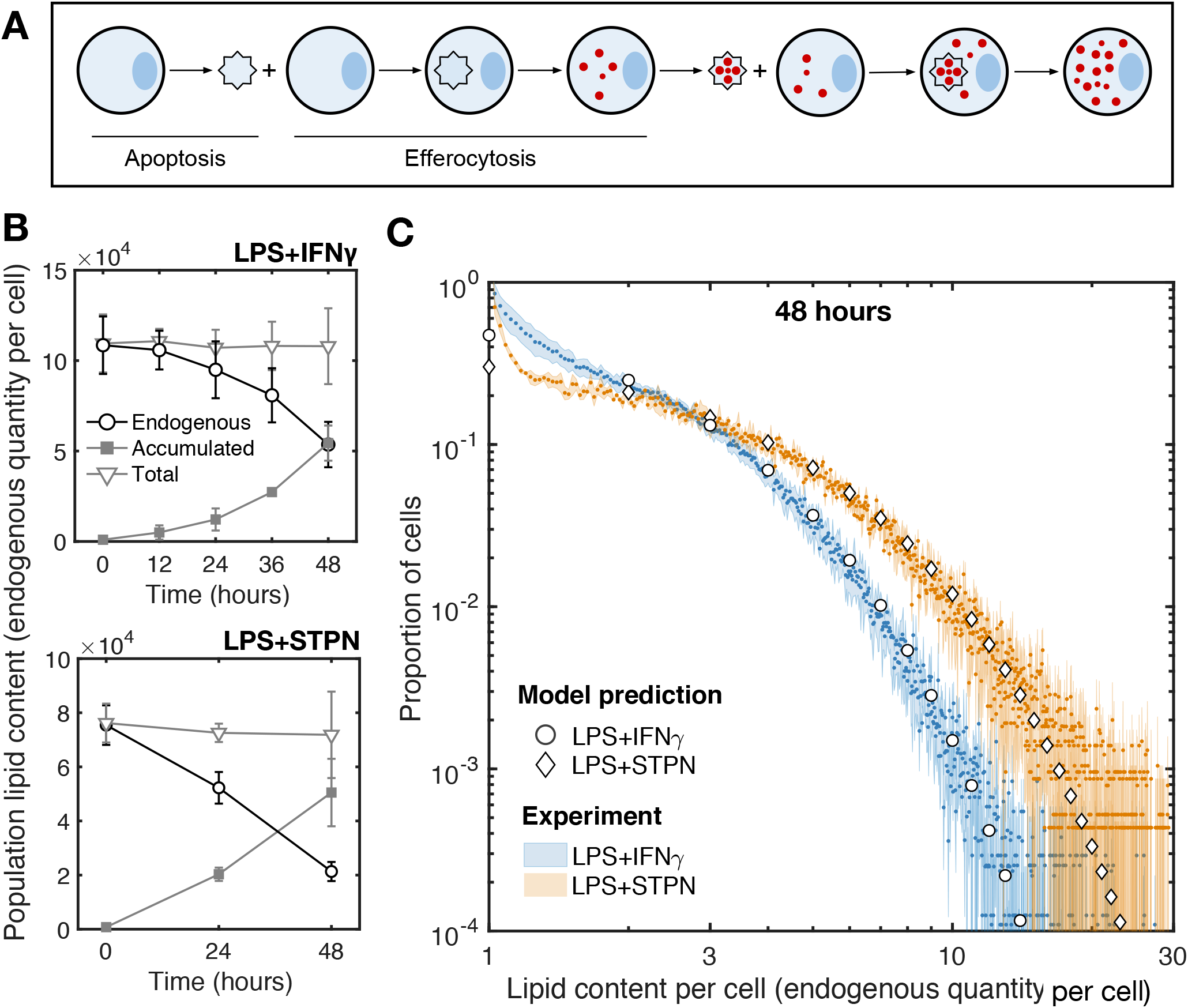
Enhanced apoptosis promotes foam cell formation. Inclusion of a dimension for lipid accumulation (in units of the endogenous quantity of lipid per cell) for model comparison. The data presented is averaged from four experiments (LPS+IFN*γ*) and three experiments (LPS+STPN) carried with different mice on different days. **(A)** Cartoon illustration of lipid droplet accumulation (red) derived from the membranes of live cells (blue circle) that have since died (blue star) and have been consumed. Here two apoptosis and efferocytosis events are depicted to show the progression of lipid accumulation. **(B)** The quantity of endogenous lipid (black circles), the total quantity of accumulated lipid (filled grey square) and their addition (light grey triangles) inside the whole BMDM population following stimulation with LPS and IFN*γ* (top) and LPS and STPN (bottom).**(C)** The proportion of cells with a range of lipid contents (in units of the quantity of endogenous lipid per cell) at 48 hours after stimulation with LPS and IFN*γ* (blue) or LPS and STPN (orange). Also shown are predictions from the mathematical model (no cell division) run to 48 hours from an initial condition where every cell contain 1 unit quantity of endogenous lipid with death rate 1/80 per hour (circles) or 1/40 per hour (diamonds).

## 3 Discussion

We hypothesised that cannibalistic efferocytosis contributes to the accumulation of substances inside inflammatory macrophages. A mathematical model (based on coagulation-fragmentation equations) was formulated and used to test this hypothesis. We concluded that the transfer of substances from dead to live macrophages via efferocytosis (Figure 1) [20, 24] continually concentrates these substances among surviving macrophages (Figure 2). Model predictions were confirmed by quantifying latex microbead and neutral lipid accumulation inside murine bone marrow-derived macrophages following their stimulation into a pro-inflammatory state (M1) with LPS and IFN*γ ex vivo* (Figures 3–5). A new in-house computer vision algorithm was used to count the number of beads and quantify stained lipid droplets per cell.

Inflammatory macrophages which initially contained small quantities of phagocytosed beads and neutral lipid (in cell membranes) accumulated large quantities of beads and lipids (in lipid droplets) per cell (Figure 3A and 4A). Neither beads nor neutral lipids are degraded during efferocytosis (Figures 1, 3B and 5A) so that, as the population reduced in size, the average number of beads or lipid droplets per cell increased (Figure 3C and 4C). Our mathematical model predicted how these substances are distributed among the surviving macrophages (Figure 3D, 4D and 5C). That is, efferocytosis caused the indigestible contents of the macrophage population to concentrate and coalesce inside the macrophage population as it declined in size. Neutral lipids that accumulated derived from the membranes of macrophages that had died and then been consumed. Consequently, enhanced macrophage cell death (induced by staurosporin) increased the rate and extent of foam cell formation (Figure 5).

Our findings suggest that any phagocytosed particle or pathogen that avoids subcellular degradation should perpetually accumulate within macrophages via efferocytosis. We experimentally demonstrated this for phagocytosed beads and endogenous neutral lipid. Consequently, we believe that this type of accumulation will occur in many other contexts. Our results suggest that efferocytosis is capable of driving the accumulation of sterile particles inside macrophages during chronic inflammation [15]. For example: (i) lung inflammation when alveolar macrophages accumulate airborne pollutants such as asbestos and silica crystals [19, 23], (ii) brain inflammation (Alzheimer’s disease) when microglia accumulate fibrilar amyloid-*β* [18] and (iii) artery wall inflammation (atherosclerosis) when monocyte-derived macrophages accumulate cholesterol crystals and neutral lipid [16]. Unlike in infections [24], cannibalistic efferocytosis has been overlooked as a mechanism of substance accumulation in these sterile inflammatory lesions. Numerous intracellular pathogens (e.g. *M. tuberculosis*) have evolved to avoid degradation and exploit efferocytosis by using dead and infected macrophages as a “Trojan horse” to infect other macrophages [20, 24, 42]. We observe this effect for beads and lipids (Figures 3 and 4) suggesting that it might also occur for sterile particles in general. That is, our results suggest that, together with pathogen replication and particle phagocytosis, the number of pathogens or sterile particles per cell can grow and diversify across macrophage populations via cannibalistic efferocytosis.

The accumulation of pathogens and sterile particles inside macrophages can induce cell death and can promote intracellular pattern-recognition receptor (PRR) activation and pro-inflammatory cytokine secretion (e.g. IL-1*β*) [12]. In this way, substance accumulation influences cell state such that macrophages might adopt similar characteristics to the macrophages which they consume. Consequently, a macrophage might become pro-inflammatory when it consumes a dead macrophage that contains a large quantity of pro-inflammatory substances. Alternatively, a macrophage might die when it consumes a dead macrophage with cytotoxic levels of pathogens or sterile particles. This “serial killing” effect is seen in *M. tuberculosis*-infected macrophages [20] and might also be caused by other substances that promote cell death, such as cholesterol [22, 43]. Since cell death and efferocytosis promote substance accumulation (Figure 5) and substance accumulation can promote cell death [20], a detrimental positive feedback loop might arise which progressively increases the rate of cell death and substance accumulation inside macrophages [44].

Macrophage foam cells commonly appear in a variety of immune responses because PRR activation (e.g. by LPS) decreases cholesterol efflux from macrophages [22, 39, 45, 46]. However, a decrease in lipid efflux from macrophages does not explain the source of lipid in macrophage foam cells. Our results suggest that foam cells arise in inflammatory macrophage populations because these cells slowly (relative to the death rate) remove accumulated lipid that is derived from the membranes of dead and consumed macrophages. Furthermore, foam cells secrete proinflammatory cytokines and are quick to die [43, 47, 48]. In this way, endogenous lipid could progressively accumulate within macrophage populations and cause macrophages to become increasingly pro-inflammatory as the immune response ages.

Similarly to lipids, efferocytosis might also amplify the intracellular quantity of other endogenous substances, such as metabolites (e.g. uric acid) and trace elements (e.g. iron), which can also be pro-inflammatory and/or cytotoxic at high levels. For example, uric acid accumulation inside macrophages promotes inflammation associated with gout [17] and iron accumulation in macrophages promotes inflammation associated with venous leg ulcers [49].

In light of our observations, we anticipate that macrophage division [5] and emigration [50] might be beneficial to the resolution of inflammation, whereas monocyte recruitment [6] might be detrimental. Macrophage division reduces substance accumulation by splitting accumulated substances between daughter cells; essentially diluting substances within macrophages (Figure 1) [35]. Macrophage emigration could remove accumulated substances from the tissue, preventing it from recycling back into the local macrophage population via efferocytosis. Lastly, monocyte recruitment introduces endogenous substances (e.g. neutral lipids) into the tissue which fuel their accumulation inside monocyte-derived macrophages via efferocytosis. We have extended the mathematical model presented here to model these type of effects of lipid accumulation inside macrophages during artery wall inflammation associated with atherosclerosis (under review) [51].

Our coalescence model generalises beyond macrophages and draws parallels between cannibalistic cell and animal populations [27, 28]. We have shown that cannibalism causes the average amount of substances per cell to exponentially increase with time (Figures 3C and 4C). In ecosystems, the amount of inorganic substances per animal can increase exponentially along the trophic levels of a food chain (i.e. biomagnification) [52]. This biomagnification process arises from the transfer of indigestible substances via predation. Thus biomagnification should also arise via cannibalism. In this way, substance accumulation via cannibalistic efferocytosis can be viewed as a biomagnification or coalescence process [31]. Intraspecific biomagnification (due to cannibalism) could be particularly detrimental if it caused the accumulation of harmful substances. For example, either inorganic toxins or infectious agents could biomagnify to harmful levels inside single cannibalistic populations. This process could contribute to increased occurrences of diseases in animal populations where the causative agent propagates via cannibalism (e.g. prion diseases Kuru neurodegenerative disorder and bovine spongiform encephalopathy) [30].

This study contributes to a growing body of evidence that casts cannibalitic efferocytosis as a double-edged sword [20, 24]. Although crucial for tissue homeostasis and inflammation resolution [25], efferocytosis also perpetuates subcellular substance accumulation which might contribute to the pathogenesis of inflammation. In this light, cannibalism has similar detrimental consequences in cell populations as in animal populations.

## 4 Methods

### 4.1 Mathematical model

#### 4.1.1 Model statement

The following system of non-linear differential equations (a coagulation-fragmentation model) was used to model the dynamic redistribution of beads inside macrophage populations via apoptosis, efferocytosis and division:

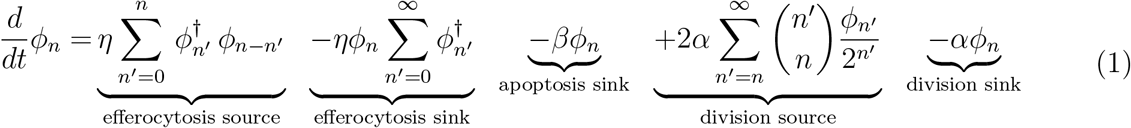

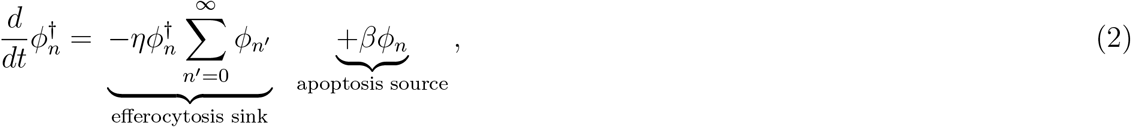

where *ϕ_n_* = *ϕ_n_*(*t*) and 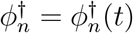 represent the number density of live and apoptotic macrophages that contain *n* ≥ 0 beads at time *t* ≥ 0. Equations (1) and (2) are closed by specifying initial distributions *ϕ_n_*(0) = *ψ_n_* and 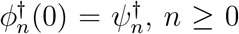. The apoptosis, efferocytosis and division rates were assumed to be independent of cellular bead content and assumed to occur with rates *β* (per unit time), *η* (per cell per unit time) and *α* (per unit time) respectively. We further assumed that bead numbers are conserved during apoptosis, efferocytosis and division. In more detail, we assume: (i) apoptosis produces one apoptotic cell with *n* beads from one live cell with *n* beads, (ii) efferocytosis transfers *n* beads contained inside one apoptotic cell (consumed) to one live cell (consumer) that contains *n*′ ≥ 0 beads to produce one live cell that contains *n* + *n*′ beads and (iii) division produces two live macrophages (daughter cells) which contain 0 ≤ *n*′ ≤ *n* and *n* − *n*′ beads from one live macrophage (parent cell) with *n* beads. The mass action kinetics are shown in Figure 2A. A convolution source term models efferocytosis when all possible bead numbers inside live and apoptotic cells are accounted for. A binomial source term models cell division when each bead inside a parent cell is equally likely to transfer to either daughter cell upon division. The total number of live and apoptotic cells are defined as 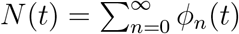 and 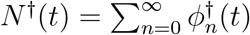 respectively.

#### 4.1.2 Model solutions

We define the proportion of live and dead cells with *n* beads by *p_n_*(*t*) ≡ *ϕ_n_*(*t*)/*N*(*t*) and 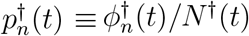 respectively. We make the simplifying assumption that efferocytosis is instantaneous (*η* → ∞) such that 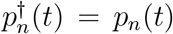. With this rescaling and assumption, equations (1) and (2) reduces to:

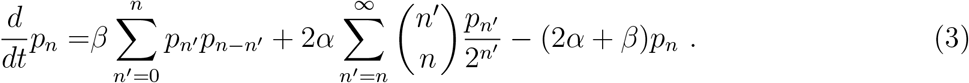

The solution for the population size is *N*(*t*) = *N*_0_*e*^(*α-β*)*t*^ where *N*_0_ is the initial number of cells.

Solutions to equation (3) are shown in Figure 2 for the case without cell division *α* = 0 (Figure 2B) and with cell division *α* = *β* (Figure 2C). Here, we use an initial condition where every cell initially (*N*_0_ cells) contains *n* =1 bead (*p_n_*(0) = 1 for *n* =1 and *p_n_*(0) = 0 for *n* ≠1). With *α* = 0, the population exponentially decays in size *N*(*t*) = *N*_0_*e*^−*βt*^ and the proportion of cells with *n* beads *p_n_* has the following geometric distribution (see Supplementary Material 1 for details):

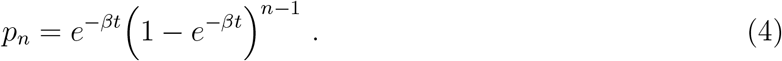

With *α* = *β* the population size remains constant over time *N*(*t*) = *N*_0_ and the proportion of cells with *n* beads *p_n_* tends to an equilibrium state that satisfies the following relation:

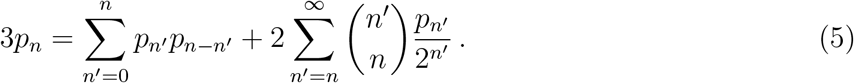

The forward Euler method [53] was used to numerically solve equation (3) with *α* = *β*.

To compare the model to the experimental data shown in Figure 3, we assumed that bead-loaded macrophages stimulated with LPS and IFN*γ* do not divide *α* = 0 and die with rate *β* = 1/60 per hour (see Supplementary Material 2). The model prediction for the proportion of cells with *n* beads (*p_n_*) at *t* = 24 and 48 hours was found by numerically solving (forward Euler method [53]) equation (3) with an initial condition *p_n_*(0) equal to the experimental initial condition.

To compare the model to the experimental data shown in Figure 5, we assumed that macrophages stimulated with LPS and IFN*γ* do not divide *α* = 0 and die with rate *β* = 1/80 per hour (see Supplementary Material 2). We also assumed that macrophages stimulated with LPS and STPN do not divide *α* = 0 and die with rate *β* = 1/40 per hour (see Supplementary Material 2). We generalise the number of beads per cell *n* in our model to also represent the lipid content in units of the endogenous neutral lipid content (i.e. in cell membranes) per cell. We assume that every cell initially contains the same single unit of endogenous lipid such that *p_n_*(0) = 1 if *n* =1 and *p_n_*(0) = 0 if *n* ≠ 1. Thus the model prediction for the proportion of cells with lipid content *n* (*p_n_*) at *t* = 24 and 48 hours was found from equation (4).

### 4.2 Experimental methods

#### 4.2.1 Cell culture

Hematopoietic stem cells were harvested from the femurbone marrow of mice. These bone marrow cells were differentiated into macrophages by culturing for 7 days at 37°C and 5% CO_2_ in 8mL high glucose Dulbecco’s modified eagle media (DMEM) supplemented with 10% head-inactivated fetal bovine serum (FBS), 1% penicillin/streptomycin (P/S) and 10% supernatent derived from L929 fibroblasts (L929-condition media) as a source of macrophage colony-stimulating factor [54] in 100mm non-tissue culture treated Petri dishes (Thermo Fisher Scientific, Sterilin, UK). On day 5, 3mL of medium was removed and an additional 5ml of medium was added. Gentle scrapping was used to lift cells off dish surface. Cells were then counted and resuspended in DMEM at the desired cell concentration.

#### 4.2.2 Bead accumulation

Cells were plated at 1 × 10^5^ cells per well of a 4-well Nun-Tek^TM^ Chamber slide™ (1.8 cm^2^) or of a 24-well plate (2 cm^2^) (Sigma-Aldrich, Gilligham, UK) with glass coverslip in high glucose DMEM supplemented with 10% FBS and 1% P/S and left at room temperature for 1 hour to adhere evenly across the plate. The cells were then incubated (37°C and 5% CO_2_) for 6 hours and then exposed to 0.5 × 10^5^ red and 0.5 × 10^5^ blue opsinised (1 hour in 10%v/v human serum) 3 μm diameter latex polystyrene beads (Sigma-Aldrich, Gilligham, UK) such that there were 1 bead per cell. Plates were centrifuged (400g for 1 min) so the beads were evenly spread on top of the macrophage population. Cells were then incubated (37°C and 5% CO_2_) for 18 hours so that most beads were phagocytosed. The medium was then replaced with high glucose DMEM supplemented with 10% FBS, 10% L929-conditioned media and 1% P/S supplemented with 100 ng/mL crude LPS (InVivogen, San Diego, CA, USA) and 20 ng/mL IFN*γ* (R&D Systems, Abingdon, UK) to polarise macrophages into an M1 state. Medium without LPS and IFN*γ* was used as a control. Cells were incubated (37°C and 5% CO_2_) for 0, 24 and 48 hours and then fixed with 4°C methanol for 3 minutes and stained with safranin (Sigma-Aldrich, Gilligham, UK) for 10 minutes. The data presented is averaged from four experiments carried with different mice on different days.

#### 4.2.3 Lipid accumulation

The protocol was the same for the bead accumulation except without the addition with beads. Cells were fixed prior to (0 hours) and 12, 24, 36, 48 and 60 hours after incubation. Cells were fixed with 4% formalin for 20 mins and stained with oil red O (Sigma-Aldrich, Gilligham, UK) for 15 mins and then stained with Mayer’s hematoxylin (Sigma-Aldrich, Gilligham, UK) for 10 mins.

#### 4.2.4 Cell quantification

The IncucyteZoom (Sartorius, Göttingen, Germany) was used to obtain the time-lapse microscopy images of bead-loaded macrophage population in 24-well plates. The Hamamatsu Nanozoomer (Hamamatsu Photonics, Japan) was used to obtain whole slide photomicrographs of fixed and stained macrophages. A new in-house image recognition algorithm was developed to quantify either the total number of beads or lipid droplets per cell for every cell in whole slide photomicrographs. The algorithm uses superpixellation and machine learning to learn from a human operator how to differentiate between areas in the image is a bead, lipid droplet, cell or background. Once trained, the algorithm then produces a classification mask across sections of whole slide photomicrographs which identifies whole areas of beads, lipid droplets, cells or background. From this classification mask, the algorithm counts the number of beads per cell (using the average area per bead) or the area of lipid droplets per cell for every cell in the image.

## 5 Acknowledgments

We thank Dan Lucy and Agata Rumianek of the University of Oxford Sir William Dunn School of Pathology for helpful discussions. Mary R. Myerscough and Hugh Z. Ford acknowledge support from an Australian Research Council Discovery Project Grant (to Mary R. Myerscough). David R. Greaves and Hugh Z. Ford acknowledge support from the British Heart Foundation (Programme Grant RG/15/10/31485 to David R. Greaves). We acknowledge the contribution to this study made by the Oxford Centre for Histopathology Research and the Oxford Radcliffe Biobank, in particular: Stephanie Jones, Emma Bowes, Becky Davies, Ying Cui and Clare Verrill.

