## Supplementary Material 1 for "Efferocytosis perpetuates substance accumulation inside macrophage populations"

### Supplementary material 1: Derivation and analysis of a coagulation-fragmentation model of substance accumulation in macrophages

#### 1 Coagulation-fragmentation model derivation

Here, we derive coagulation-fragmentation equations that model the accumulation of beads (and other indigestible substances) inside macrophage populations.

##### 1.1 Model statement

Let continuous dependent variables $\phi_n(t) \geq 0$ and $\phi_n^\dagger(t) \geq 0$ respectively represent the number density of live and dead macrophages with $n \geq 0$ beads (discrete independent variable) at time $t \geq 0$ (continuous independent variable). Let $\phi(t) \equiv \{\phi_1(t),\, \phi_2(t),\, \ldots\}$ and $\phi^\dagger(t) \equiv \{\phi_1^\dagger(t),\, \phi_2^\dagger(t),\, \ldots\}$. We model the time evolution of $\phi_n(t)$ and $\phi_n^\dagger(t)$ via the following system of coupled non-linear ordinary differential equations:

$$\frac{d}{dt}\phi_n(t) = D_n - D_n' - A_n + E_n - E_n'\,, \tag{1}$$

$$\frac{d}{dt}\phi_n^\dagger(t) = A_n' - E_n''\,, \tag{2}$$

where the positive functions:

- $D_n = D_n(\phi(t), \phi^\dagger(t))$ and $D_n' = D_n'(\phi(t), \phi^\dagger(t))$ respectively represent the source and sink term associated with division for live cells that contain $n$ beads,
- $A_n = A_n(\phi(t), \phi^\dagger(t))$ and $A_n' = A_n'(\phi(t), \phi^\dagger(t))$ respectively represent the sink and source term associated with apoptosis for live and dead cells that contain $n$ beads,
- $E_n = E_n(\phi(t), \phi^\dagger(t))$ and $E_n'(\phi(t), \phi^\dagger(t))$ respectively represent the source and sink term associated with efferocytosis for live cells that $n$ beads and
- $E_n'' = E_n''(\phi(t), \phi^\dagger(t))$ represents the sink term associated with efferocytosis for dead cells that contain $n$ beads.

The total numbers of live $N(t)$ and dead $N^\dagger(t)$ cells are given by:

$$N(t) \equiv \sum_{n=0}^{\infty} \phi_n(t) \quad \text{and} \quad N^\dagger(t) \equiv \sum_{n=0}^{\infty} \phi_n^\dagger(t)\,. \tag{3}$$

We denote the initial number of live and dead cells as $N(0) = N_0$ and $N^\dagger(0) = N_0^\dagger$ respectively. We close equations (1) and (2) by assuming that $\Phi_n$ live and $\Phi_n^\dagger$ dead cells initially contain $n$ beads such that:

$$\phi_n(0) = \Phi_n \text{ and } \phi_n^\dagger(0) = \Phi_n^\dagger\,. \tag{4}$$

#### 1.2 Specifying the functional forms

The functional forms for $A$, $A'$, $E$, $E'$, $E''$, $D$ and $D'$ that appear in equations (1) and (2) are now introduced.

##### 1.2.1 Apoptosis

We assume that apoptosis: (i) occurs at a constant rate $\beta$ per unit time, (ii) is independent of bead content and (iii) conserves bead numbers. So in equations (1) and (2) apoptosis is modelled by the following equal and opposite sink and source terms:

$$A_n = A_n' = \beta \phi_n(t)\,. \tag{5}$$

##### 1.2.2 Efferocytosis

We assume that efferocytosis (i) occurs at a constant rate $\eta$ per cell per unit time, (ii) is independent of bead content and (iii) conserves bead numbers. So in equation (2) efferocytosis is modelled by a sink term for the number of dead cells which is equal to the rate at which dead cells containing $n$ beads are consumed by any live cell:

$$E'' = \eta \phi_n^{\dagger}(t) \sum_{n'=0}^{\infty} \phi_{n'}(t) = \eta \phi_n^{\dagger}(t) N(t)\,. \tag{6}$$

In equation (1) we account for efferocytosis via two terms. A sink term $E'$ that represents the rate at which live cells containing $n$ beads consume dead cells containing $n' = 1, 2, \ldots$ beads:

$$E' = \eta \phi_n(t) N^{\dagger}(t)\,. \tag{7}$$

The source term $E$ is modelled by a discrete convolution which enumerates all the ways in which a live cell containing $n$ beads can be produced when a live cell containing $n' = 0, 1, \ldots, n$ beads consumes a dead cell containing $'n - n'$ beads:

$$E = \eta \sum_{n'=0}^{n} \phi_{n'}(t) \phi_{n-n'}^{\dagger}(t)\,. \tag{8}$$

##### 1.2.3 Division

We assume that division: (i) occurs at a constant rate $\alpha$ per unit time, (ii) is independent of bead content, (iii) splits the number of beads contained by the parent cell between two daughter cells and (iv) We each bead inside the parent cell is equally likely to end up in either daughter cell. In equation (1) we account for cell division via two terms A sink term $D'$ that represents the rate at which cells containing $n$ beads divide:

$$D' = \alpha \phi_n(t)\,, \tag{9}$$

The source term $D$ is modelled by the binomial distribution that enumerates the the likelihood that a parent cell with $n' = n, n+1, \ldots$ beads produces daughter cells that contain $n$ beads and $n' - n$ beads:

$$D = 2\alpha \sum_{n'=n}^{\infty} \frac{1}{2^{n'}} \binom{n'}{n} \phi_{n'}(t)\,. \tag{10}$$

##### 1.3 The full dimensional model

Substituting equations (5)-(10) into equations (1) and (2) produces the following coagulation-fragmentation equations for $\phi_n(t)$ and $\phi^\dagger_n(t)$:

$$\frac{d}{dt}\phi_n(t) = \underbrace{\eta \sum_{n'=0}^{n} \phi^\dagger_{n'}(t)\,\phi_{n-n'}(t)}_{\text{efferocytosis source}} \underbrace{-\eta\phi_n(t)\sum_{n'=0}^{\infty}\phi^\dagger_{n'}(t)}_{\text{efferocytosis sink}} + \underbrace{\alpha\sum_{n'=n}^{\infty}\binom{n'}{n}\frac{\phi_{n'}(t)}{2^{n'-1}}}_{\text{division source}} \underbrace{-\alpha\phi_n(t)}_{\text{division sink}} \underbrace{-\beta\phi_n(t)}_{\text{apoptosis sink}} \tag{11}$$

$$\frac{d}{dt}\phi^\dagger_n(t) = \underbrace{\beta\phi_n(t)}_{\text{apoptosis source}} \underbrace{-\eta\phi^\dagger_n(t)\sum_{n'=0}^{\infty}\phi_{n'}(t)}_{\text{efferocytosis sink}}\ . \tag{12}$$

The initial condition is given by equation (4).

##### 1.4 Two differently coloured beads

In our experiments we also considered two differently coloured beads (red and blue). It is straightforward to extend equations (11) and (12) to account for differently coloured beads.

Let $\psi_{i,j}(t)$ and $\psi^\dagger_{i,j}(t)$ resepctively represent the number of live and dead cells that contain $i \geq 0$ red beads and $j \geq 0$ blue beads at time $t$. As such, $\phi_n(t)$ represents the number of cells that contain a total of $n = i + j$ beads (these may be red or blue) so that:

$$\phi_n(t) = \sum_{i+j=n}\psi_{i,j}(t) = \sum_{n'=0}^{n}\psi_{n',n-n'}(t)\,. \tag{13}$$

As for the case with one bead type, the kinetics of apoptosis, efferocytosis and division are given by:

$$\begin{aligned}
\psi_{i,j} &\xrightarrow[\beta]{\text{apoptosis}} \psi^\dagger_{i,j}\\
\psi^\dagger_{i,j} + \psi_{i',j'} &\xrightarrow[\eta]{\text{efferocytosis}} \psi_{i+i',j+j'}\\
\psi_{i,j} &\xrightarrow[\alpha]{\text{division}} \psi_{i',j'} + \psi_{i-i',j-j'}\ ,
\end{aligned}$$

for some $i \geq i' \geq 0$ and $j \geq j' \geq 0$. We assume that the accumulation of differently coloured beads are independent. Using the same arguments made in section 1.2, we deduce that the time-evolution of $\psi_{i,j}(t)$ and $\psi^\dagger_{i,j}(t)$ are given by the following pair of coagulation-fragmentation equations:

$$\frac{d}{dt}\psi_{i,j} = \eta\sum_{i'=0}^{i}\sum_{j'=0}^{j}\psi^\dagger_{i',j'}\psi_{i-i',j-j'} - \eta\psi_{i,j}\sum_{i'=0}^{\infty}\sum_{j'=0}^{\infty}\psi^\dagger_{i',j'} + \alpha\sum_{i'=i}^{\infty}\sum_{j'=j}^{\infty}\binom{i'}{i}\binom{j'}{j}\frac{\psi_{i',j'}}{2^{i'+j'-1}} - (\alpha+\beta)\psi_{i,j}\ , \tag{14}$$

$$\frac{d}{dt}\psi^\dagger_{i,j} = \beta\psi_{i,j} - \eta\psi^\dagger_{i,j}\sum_{i'=0}^{\infty}\sum_{j'=0}^{\infty}\psi_{i',j'}\ . \tag{15}$$

We close equations (14) and (15) by impoing the following initial condition:

$$\psi_{i,j}(0) = \Psi_{i,j} \quad \text{and} \quad \psi^\dagger_{i,j}(0) = \Psi^\dagger_{i,j}\ . \tag{16}$$

#### 1.5 Nondimensionalisation

Here we nondimensionalise equations (11) and (12) and equations (14) and (15). The number density of live and dead cells with $n$ beads are scaled with the total number of live and dead cells respectively so that:

$$p_n(t) \equiv \phi_n(t)/N(t) \;\text{ and }\; p^\dagger_n(t) \equiv \phi^\dagger_n(t)/N^\dagger(t)\,, \tag{17}$$

where $p_n(t)$ and $p^\dagger_n(t)$ are the population density of live and dead cells respectively:

$$\sum_{n=0}^{\infty} p_n(t) = \sum_{n=0}^{\infty} p^\dagger_n(t) = 1\,. \tag{18}$$

Similarly, we let:

$$q_n(t) \equiv \psi_{i,j}(t)/N(t) \;\text{ and }\; q^\dagger_n(t) \equiv \psi^\dagger_{i,j}(t)/N^\dagger(t)\,. \tag{19}$$

The total number of live and dead cells are nondimesnionalised by dividing both by the initial number of live $N_0$ cells:

$$\widetilde{N}(t) \equiv N(t)/N_0 \text{ and } \widetilde{N}^\dagger(t) \equiv N^\dagger(t)/N_0. \tag{20}$$

Time is rescaled with $\beta^{-1}$, the mean macrophage lifetime, so that:

$$\widetilde{t} \equiv \beta t\,. \tag{21}$$

Substituting equations (17)-(21) into equations (11), (12) and (4) produces the following system of ordinary differential equations for the time-evolution of the proportion of live and dead cells with $n$ beads:

$$\frac{d}{d\widetilde{t}}p_n + \frac{p_n}{\widetilde{N}}\frac{d}{dt}\widetilde{N} = b\widetilde{N}^\dagger\Big(\sum_{n'=0}^{n} p^\dagger_{n'}p_{n-n'} - p_n\Big) + 2a\sum_{n'=n}^{\infty}\binom{n'}{n}\frac{p_{n'}}{2^{n'}} - (1+a)p_n \tag{22}$$

$$\frac{d}{d\widetilde{t}}p^\dagger_n + \frac{p^\dagger_n}{\widetilde{N}^\dagger}\frac{d}{d\widetilde{t}}\widetilde{N}^\dagger = \frac{\widetilde{N}}{\widetilde{N}^\dagger}\Big(p_n - b\widetilde{N}^\dagger p^\dagger_n\Big)\,, \tag{23}$$

with:

$$p_n(0) = \frac{\Phi_n}{N_0} \text{ and } p^\dagger_n(0) = \frac{\Phi^\dagger_n}{N^\dagger_0}\,, \tag{24}$$

and:

$$a = \frac{\alpha}{\beta} \;\text{ and }\; b = \frac{\eta N_0}{\beta}\,. \tag{25}$$

Parameter $a$ represents the rate of division ($\alpha$) relative to the rate of death ($\beta$). Parameter $b$ represents the initial rate of efferocytosis ($\eta N_0$) relative to the rate of death. Hereafter, we neglect the tilde notation.

#### 1.6 Solution for population size

In order to determine expressions for $N(t)$ and $N^{\dagger}(t)$, we note from equation (18) that $\sum_{n=0}^{\infty}\sum_{n'=n}^{\infty}\binom{n'}{n}\frac{p_{n'}(t)}{2^{n'}} = 1$ and $\sum_{n=0}^{\infty}\sum_{n'=0}^{n} p_{n'}(t)p_{n-n'}(t) = 1$. Thus summing equations (22) and (23) across all $n$ yields the following ordinary differential equations for $N(t)$ and $N^{\dagger}(t)$ respectively:

$$\frac{d}{dt}N = (a-1)N \ , \quad N(0) = 1 \quad \text{and} \quad \frac{d}{dt}N^{\dagger} = N\Big(1 - bN^{\dagger}\Big) \ , \quad N^{\dagger}(0) = \frac{N_0^{\dagger}}{N_0} \ . \tag{26}$$

which have solutions:

$$N = e^{(a-1)t} \quad \text{and} \quad N^{\dagger} = \frac{1}{b} + \Big(\frac{N_0^{\dagger}}{N_0} - \frac{1}{b}\Big)e^{\frac{b}{a-1}(1-e^{(a-1)t})} \ . \tag{27}$$

As $t \to \infty$ the number of live cells either grows $N(t) \to \infty$, decays $N(t) \to 0$ or remains steady $N(t) = N_0$ when $a > 1$, $a < 1$ and $a = 1$ respectively. As time $t \to \infty$ the number of dead cells either tends to the finite number $\frac{1}{b}$, $\frac{1}{b} + \Big(\frac{N_0}{N_0^{\dagger}} - \frac{1}{b}\Big)e^{\frac{b}{a-1}}$ or $\frac{N_0}{N_0^{\dagger}}$ when $a > 1$, $a < 1$ and $a = 1$ respectively.

For simplicity, we assume that the number of dead cells is initially at the steady state value such that it remains constant over time:

$$N_0^{\dagger} = \frac{N_0}{b} = \frac{\beta}{\eta} \implies N^{\dagger}(t) = N^{\dagger}(0) = \frac{N_0^{\dagger}}{N_0} = \frac{1}{b} \text{ for all } t \, , \ a \text{ and } b > 0 \ . \tag{28}$$

#### 1.7 The dimensionless model

Substituting equations (27) and (**??**) into equations (22) and (23) produces the following coagulation-fragmentation equations:

$$\frac{d}{dt}p_n = \sum_{n'=0}^{n} p_{n'}^{\dagger}p_{n-n'} + 2a\sum_{n'=n}^{\infty}\binom{n'}{n}\frac{p_{n'}}{2^{n'}} - (2a+1)p_n \tag{29}$$

$$\frac{d}{dt}p_n^{\dagger} = be^{(a-1)t}\Big(p_n - p_n^{\dagger}\Big) \ . \tag{30}$$

Similarly, the case with two differently coloured beads can be modelled by the following 2D coagulation-fragmentation equation:

$$\frac{d}{dt}q_{i,j} = \sum_{i'=0}^{j}\sum_{j'=0}^{j} q_{i',j'}^{\dagger}q_{i-i',j-j'} + 2a\sum_{i'=i}^{\infty}\sum_{j'=j}^{\infty}\binom{i'}{i}\binom{j'}{j}\frac{q_{i',j'}}{2^{i'+j'}} - (1+2a)q_{i,j} \ , \tag{31}$$

$$\frac{d}{dt}q_{i,j}^{\dagger} = be^{(a-1)t}\Big(q_{i,j} - q_{i,j}^{\dagger}\Big) \ . \tag{32}$$

#### 1.8 Analytically tractable model

In general, it is not possible to construct explicit analytical solutions to equations (29) and (30) or equations (31) and (32). However the time-dependent solution can be found analytically for the special case when $a = 0$ (no cell division), $b \to \infty$ (instant apoptotic cell consumption) and $\Phi_n = N_0\delta(n-1)$ and $\Psi_{i,j} = \frac{N_0}{2}\Big(\delta(j-1)\delta(i) + \delta(j)\delta(i-1)\Big)$ where $\delta$ is the Dirac delta function such that $\delta(x) = 1$ if $x = 0$ and $\delta(x) = 0$ (every cell initially contains one bead, half of which are red and the other half blue). From equations (27) and (30), the assumption $b \to \infty$ implies that no apoptotic cells exist $N^{\dagger} = 0$ and the proportion of live and dead cells with $n$ beads are equal $p_n^\dagger = p_n$. Thus, under these assumptions, equations (29) and (30) simplify to

$$\frac{d}{dt}p_n = \sum_{n'=1}^{n-1} p_{n'}p_{n-n'} - p_n \ , \quad p_n(0) = \delta(n-1) \ , \tag{33}$$

and equations (31) and (32) simplify to

$$\frac{d}{dt}q_{i,j} = \sum_{i'=0}^{i}\sum_{j'=0}^{j} q_{i',j'}q_{i-i',j-j'} - q_{i,j} \ , \quad q_{i,j}(0) = \frac{1}{2}\delta(i-1)\delta(j) + \frac{1}{2}\delta(i)\delta(j-1) \ , \tag{34}$$

Equations (33) and (34) are 1D and 2D Smoluchowski coagulation equations. Note that in this model, the total population size (dimensional) is $N(t) = N_0 e^{-t}$ and the total number of beads inside all cells is $n_{\text{tot}} = N_0$. Therefore the average number of beads per cell exponentially grows with time $n_{\text{avg}}(t) = n_{\text{tot}}/N(t) = e^t$.

### 2 Analytical solutions (no cell division)

#### 2.1 1D case: total number of (red and blue) beads per cell

Here we determine the time dependent solution to equation (33). We use a generating function approach used to solve Smoluchowski coagulation equations [1–3]. First, we define:

$$f(z,t) \equiv \sum_{n=1}^{\infty} p_n(t)e^{-nz} \ , \tag{35}$$

noting from equation (33) that:

$$f(z,0) = e^{-z} \ . \tag{36}$$

By the Cauchy product of $f(z,t)$, we have that:

$$f^2(z,t) = \sum_{n=2}^{\infty}\sum_{n'=1}^{n} p_{n'}(t)p_{n-n'}(t)e^{-nz} \, . \tag{37}$$

Using equations (35)-(37), equation (33) supplies the following partial differential equation for $f(z,t)$;

$$\frac{\partial}{\partial t}f = \sum_{n=1}^{\infty}\frac{dp_n}{dt}e^{-nz} = \sum_{n=2}^{\infty}\sum_{n'=1}^{n-1} p_{n'}p_{n-n'}e^{-nz} - \sum_{n=1}^{\infty} p_n e^{-nz} = f^2 - f \, . \tag{38}$$

The solution to equation (38) can be found by (i) integrating with respect to $t$, (ii) applying the initial condition shown in equation (36) and (iii) using the Maclaurin series expansion $\sum_{n=0}^{\infty}(ax)^n = (1-ax)^{-1}$:

$$f(z,t) = e^{-z-t}\Big(1-(1-e^{-t})e^{-z}\Big)^{-1} = e^{-t}\sum_{n=1}^{\infty}\Big(1-e^{-t}\Big)^{n-1}e^{-nz} \, . \tag{39}$$

Comparison of equations (35) and (39) then supplies the solution for $p_n(t)$:

$$p_n(t) = e^{-t}\Big(1-e^{-t}\Big)^{n-1} \, . \tag{40}$$

As sketch of $p_n(t)$ is shown in Figure 1. We can rewrite equation (40) in terms of $N$ and $N_0$ using equation (27). Substituting $t = -\log(N/N_0)$ into equation (40) produces the following geometric distribution:

$$p_n\Big(\frac{N}{N_0}\Big) = \frac{N}{N_0}\Big(1 - \frac{N}{N_0}\Big)^{n-1} . \tag{41}$$

This expression specifies the proportion of cells with $n$ beads when the population size has decreased from $N$ to $N_0$. It is useful for comparison with experimental data.

We can determine the maximum value of $p_n(t)$ for a particular value of $n$ and the time at which the maximum occurs. By setting the first derivative of equation (40) to zero supplies:

$$\frac{d}{dt}p_n(t) = -e^{-t}\Big(1 - e^{-t}\Big)^{n-1} + (n-1)e^{-2t}\Big(1 - e^{-t}\Big)^{n-2} = 0 \implies t = \log(n) . \tag{42}$$

Note that this value of $n$ coincides with the average number of beads per cell $n_{\text{avg}} = e^t$. The maximum value of $p_n(t)$ is given by:

$$\max p_n(t) = p_n\big(\log(n)\big) = \frac{(n-1)^{n-1}}{n^n} = \frac{1}{n}\Big(1 - \frac{1}{n}\Big)^{n-1} \to e^{-1}n^{-1} \text{ as } n \to \infty . \tag{43}$$

#### 2.2 Generalisation for two types of beads

We solve equation (34) using the same approach as when there is only one type of bead. In more detail, we introduce:

$$g(x,y,t) \equiv \sum_{i=0}^{\infty}\sum_{j=0}^{\infty} q_{i,j}(t)e^{-ix-jy} . \tag{44}$$

noting from equation (34) that:

$$g(x,y,0) = \frac{1}{2}\big(e^{-x} + e^{-y}\big) . \tag{45}$$

We note, by the 2D Cauchy product of $g(x,y,t)$, that:

$$g^2 = \sum_{r=0}^{\infty}\sum_{i=0}^{\infty}\sum_{j'=0}^{j}\sum_{i'=0}^{i} q_{i',j'}q_{i-i',j-j'}e^{-ix-jy} . \tag{46}$$

Using equations (44)-(46), we can convert equation (34) into the following partial differential equation:

$$\frac{\partial}{\partial t}g = g^2 - g . \tag{47}$$

The solution to equation (47) can be found by (i) integrating with respect to $t$, (ii) applying the initial condition shown in equation (45) and (iii) using the Maclaurin series expansion:

$$g(x,y,t) = \frac{1}{2}e^{-t}(e^{-x} + e^{-y})\Big(1 - \frac{1}{2}(1 - e^{-t})(e^{-x} + e^{-y})\Big)^{-1} \tag{48}$$

$$= e^{-t}\sum_{k=0}^{\infty}\sum_{\ell=0}^{k+1} \frac{1}{2^{k+1}}(1 - e^{-t})^k\binom{k+1}{\ell}e^{-\ell x-(k+1-\ell)y} . \tag{49}$$

With $i = \ell$ and $j = k+1-\ell$, comparison of equations (44) and (49) reveals that the solution for $q_{i,j}(t)$ is:

$$q_{i,j}(t) = \frac{1}{2^{i+j}}\binom{i+j}{i}e^{-t}(1-e^{-t})^{i+j-1}\,,\ q_{0,0}(t) = 0\,. \tag{50}$$

We remark that this expression is equal to the proportion of cells with $n = i + j$ beads $p_{i+j}(t)$ (equation (40)) multiplied by the binomial distribution:

$$q_{i,j} = \frac{1}{2^{i+j}}\binom{i+j}{i}p_{i+j} = \frac{1}{2^{i+j}}\binom{i+j}{j}p_{i+j}\,. \tag{51}$$

The binomial distribution $\frac{1}{2^{i+j}}\binom{i+j}{i} = \frac{1}{2^{i+j}}\binom{i+j}{j}$ gives the likelihood of observing $i$ red and $j$ blue beads inside a cell that contains $n = i+j$ beads. Therefore the solution $q_{i,j}(t)$ is analogous to the solution $p_n(t)$ for $n = i + j$ $(n \geq i, j \geq 0)$.

##### 2.2.1 Solution description

Figure 1 displays the analytical solution for $p_n$, the proportion of cells with $n \geq 1$ beads, given by equation (41) and $q_{i,j}$, the proportion of cells with $i \geq 0$ red and $j \geq 0$ blue $(n = i+j \geq 1)$ beads, given by equation (51). As the population size declines, the population density $p_n$ becomes approximately uniform across a range of $n$ values that grows from $n = 1$.

When a bead content coincides with the average bead content $n = n_{\text{avg}} = e^t$ we have that $p_{n_{\text{avg}}} \approx n_{\text{avg}}^{-1}e^{-1} = e^{-(t+1)}$ (equation (43)) and $p_1 = e^{-t} = n_{\text{avg}}^{-1}$ (equation (40)). Thus the maximum difference in the solution across the domain $1 \leq n \leq n_{\text{avg}}$ exponentially decreases over time $p_1 - p_{n_{\text{avg}}} \approx (e-1)e^{-(t+1)}$. Therefore the population exponentially decays in size approximately as a uniform distribution ($p_n \approx p_1 = e^{-t} = \frac{N}{N_0}$) across an exponentially growing range of number of beads per cell ($1 \leq n < e^t = \frac{N_0}{N}$). That is, the average number of beads per cell and the extent of bead accumulation increases proportional to the decay in population size. And the population heterogeneity (or cell-to-cell variation) with respect to the number of beads per cell also increases as cell numbers decrease.

Furthermore, the population density remains binomially distributed across the number of red $i$ and blue $j = n-i$ beads per cell. Since the number of red $i$ and blue $n = j-i$ beads are distributed binomially, the proportion of red and blue beads per cell become more-or-less equal in cells that accumulate large numbers of beads $n \gg 1$. For example, macrophages with $i = 50$ red and $j = 50$ blue beads are far more likely ($\approx 8\%$) to be observed that macrophages with $i = 75$ red and $j = 25$ blue beads ($\approx 0\%$).

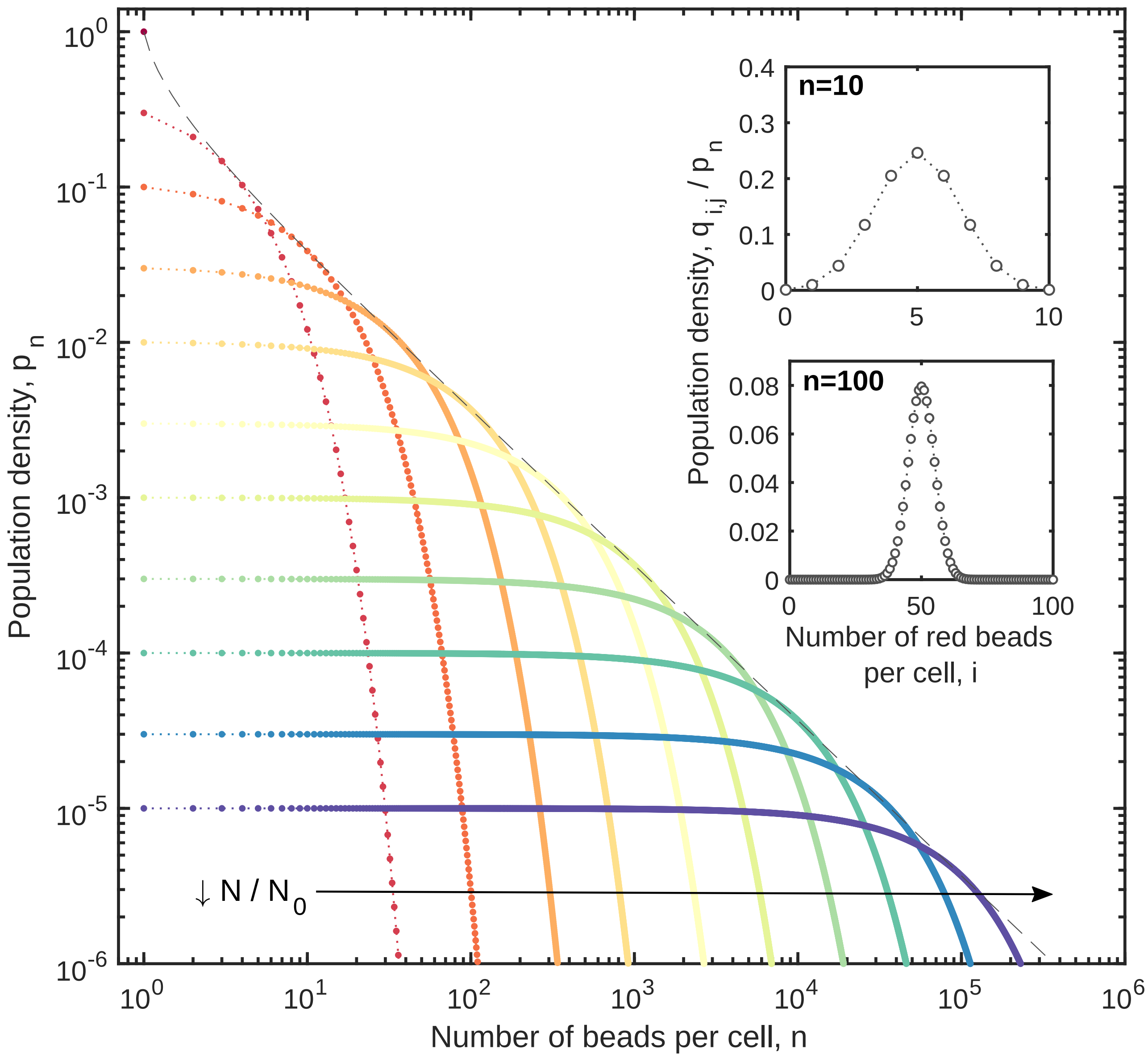


Figure 1: The evolution of the population density distribution across the total number of beads per cell $n$, $p_n$ (equation (41)), and across the number of red beads per cell $i$ and blue beads per cell $j = n - i$, $q_{i,j}$ (equation (51)), as the population size decays $N/N_0 \to 0$. Equation (41) is the analytical solution to equation (33) and equation (51) is the analytical solution to equation (34). Each coloured line represents solution $p_n$ across $n$ when the population size is $N/N_0 = 10^0, 3 \times 10^{-1}, 10^{-1}, 3 \times 10^{-2}, \ldots, 10^{-5}$ from red to blue with the envelope function for $p_n$, $n^{-n}(n-1)^{n-1}$ (dotted grey). The inset plots display the solution for the proportion of the cells with $n$ beads that contain $i$ red and $j = n - i$ blue beads $q_{i,n-i}/p_n = \frac{1}{2^n}\binom{n}{i}$ (binomial distribution) for the case where $n = 10$ (top) and $n = 100$ (bottom).

#### 3 Case with cell division

We now consider the case with cell division and perfect efferocytosis ($a > 0$ and $b \to \infty$). Numerical solutions to equations (29) and (30) were generated using the forward Euler scheme [4]. Typical results are presented in Figure 2 for cases with: (i) no cell division ($a = 0$), (ii) a cell division rate half of the apoptosis rate ($a = 0.5$), (iii) a cell division rate equal to the apoptosis rate ($a = 1$) and (iv) a cell division

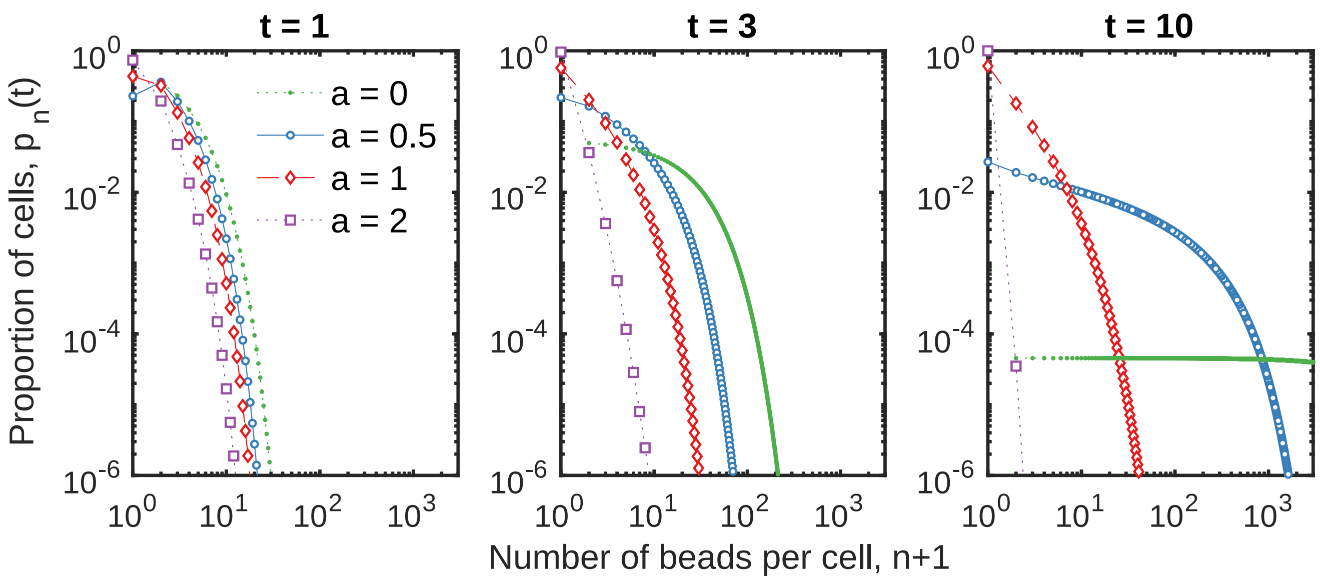


Figure 2: Numerical simulations for time evolution of the proportion of live cells with $n$ beads $p_n(t)$ given by equation (29) for $a = 0$ (green filled dots), $a = 0.5$ (blue asterisks), $a = 1$ (red open circle) and $a = 2$ (purple square) at times $t = 1$, 3 and 10 (from top-to-bottom row).

rate double the apoptosis rate ($a = 2$).

Figure 2 shows that the dilution of beads via cell division opposes the concentration of beads via apoptosis/efferocytosis. In this way, cell division can stunt, halt or reverse bead accumulation via efferocytosis. In our model the number of beads per cell is conserved during cell division, death and efferocytosis such population growth or decay respectively dilutes and concentrates beads inside the population. However, any amount of division ($a > 0$) substantially lowers the extent of bead accumulation in the population compared to the case without cell division ($a = 0$).

When cell division and apoptosis rates are balanced ($a = 1$) macrophages numbers remain constant [5, 6] and the population density tends to a steady distribution. In this state, the beads dynamically redistribute within the cell population while the expected number of cells with $n$ beads $p_n(t)$ remains constant over time. Thus equal division and apoptosis rates produce stability in both the number of macrophages (via balanced cell source and sink effects) and the numbers of beads per cell (via balanced dilution and concentration effects).

The steady state population density distribution is given by the following system of algebraic equations found by setting $a = 1$, $\frac{d}{dt}p_n = 0$ and $\frac{d}{dt}p_n^{\dagger} = 0$ in equations (29) and (30):

$$3p_n = \sum_{n'=0}^{n} p_{n'}p_{n-n'} + 2\sum_{n'=n}^{\infty} \binom{n'}{n} \frac{p_{n'}}{2^{n'}} . \tag{52}$$

There are an infinite number of solutions to equation (52). The particular solution is set by the total number of beads per cell, given by $\sum_{n\geq 0} np_n$. Solutions to equation (52) can be found using MATLABs nonlinear equation solver. Figure 3 displays these solutions when the average number of beads per cell is $\sum_{n\geq 0} np_n = 1$, 10 and 100. From Figure 3 we see that an increase in the average number of beads per cell increases the extent of bead accumulation inside macrophages. This implies that when the total number

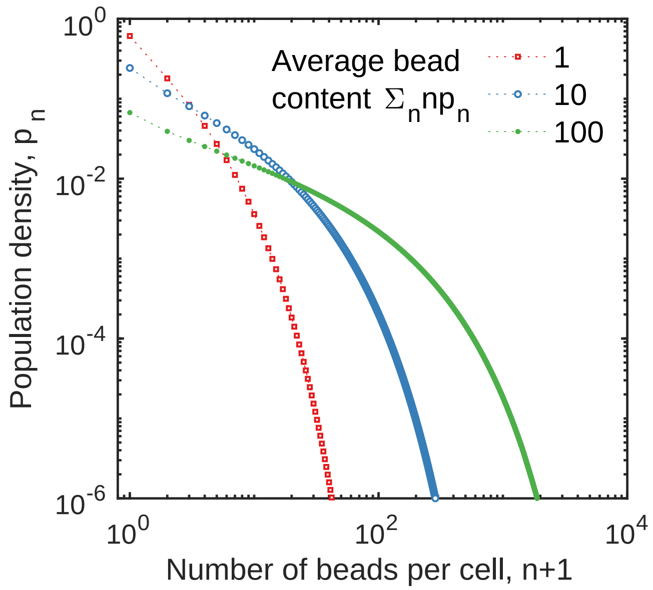


Figure 3: Numerical solutions to equation (52) showing the long time proportion of cells with $n$ beads $p_N$ when there are on average $\sum_{n\geq 0} np_N = 1$ (red squares), 10 (blue open circles) and 100 (green filled circles) beads per cell.

of beads inside the population remains fixed,the population can grow in size to obtain a new steady state distribution with a smaller extent of bead accumulation.

In this light, cell division can be viewed as a beneficial mechanism that spreads, maintains and dilutes harmful particles inside macrophage populations.
