## Supplementary Material 2 for "Efferocytosis perpetuates substance accumulation inside macrophage populations"

### Supplementary material 2: Cell quantification

#### 1 Comparison between automatic count by our algorithm and manual counting by human operators

To validate our in-house vision algorithm, an image of bead-loaded macrophages was chosen to be counted (the number of beads per cell) manually by two human operators and automatically by the algorithm. For this we selected one section of a whole slide photo (3%) of the bead-loaded macrophage population 2 days after stimulation with LPS and IFN$\gamma$ (Figure 3 in the main text). A small section of the selected image is displayed in Figure 1. Also shown is the classification mask over the selected image generated by the algorithm. This mask depicts the areas which the algorithm identified as a bead or as a cell. Following specification of the average number of pixels per bead, the algorithm then approximates the number of beads per cell from the number of pixels per cell that it identifies as a bead.

Figure 2A shows the number of cells with $n$ beads counted by the algorithm and both human operators. We see that the number of cells counted with $n$ beads are qualitatively similar for each counting approaches. The variation in numbers counted by each operator is qualitatively similar to the variation in numbers counted by the algorithm. Overall, the total number of cells counted by the algorithm is less

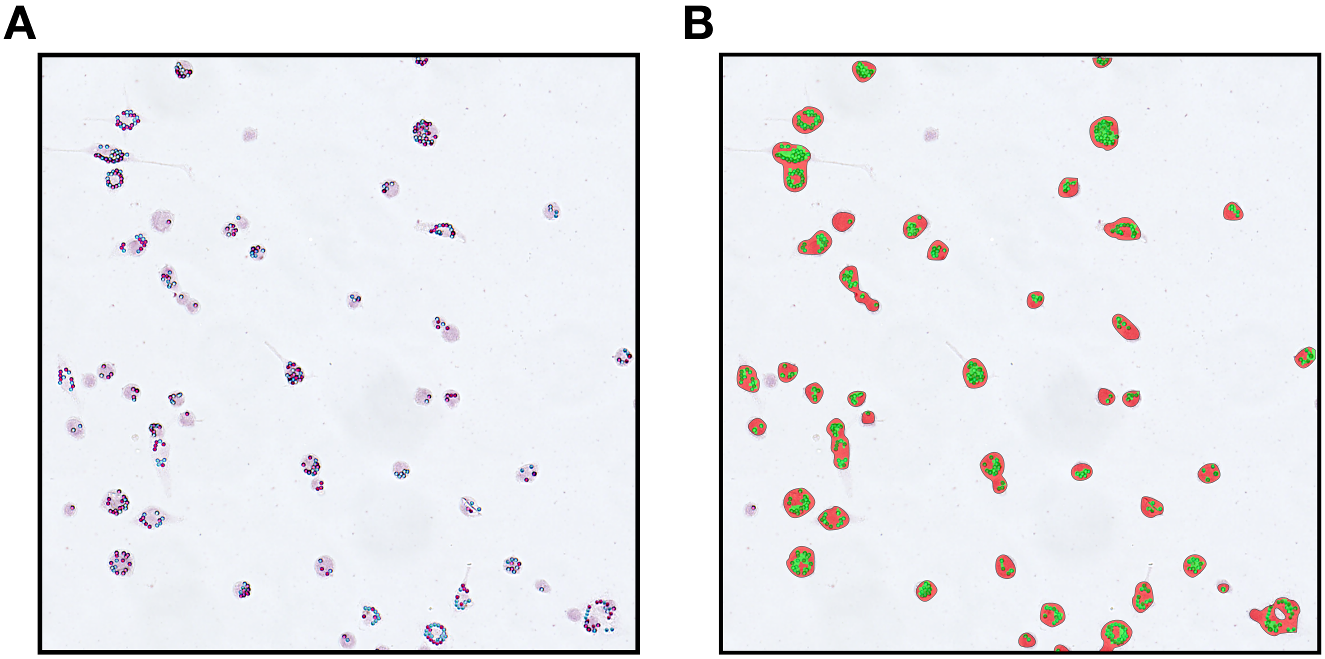


Figure 1: **The algorithm classification mask generated over a representive image.** (**A**) A representative section of the image used to count the number of beads per cell manually by 2 human operators and automatically using the algorithm. (**B**) The classification mask that depicts the areas identified as bead (green) or as cell (red) by the algorithm.

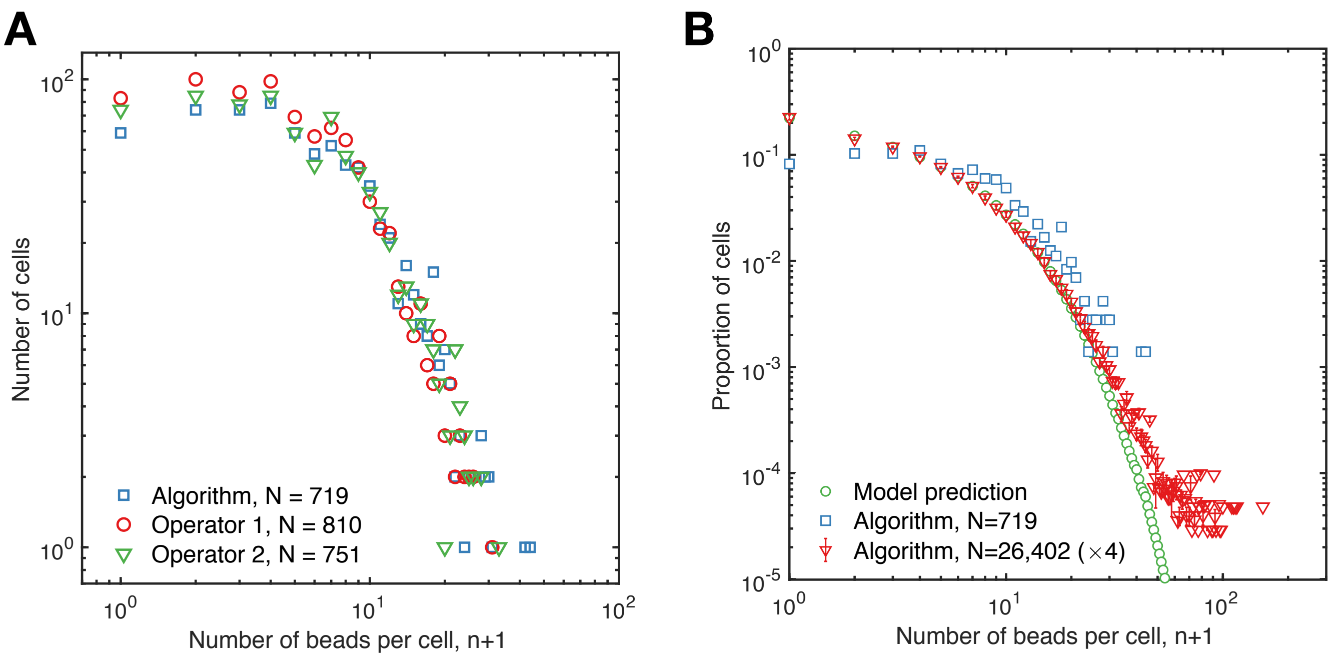


Figure 2: **Comparison between manual and automatic counting (A)** The number of cells with $n$ beads counted by the algorithm (blue squared) and both human operators (red circles and green traingles) in a representative image (a proportion of which is shown in Figure 1). The algorithm counted 719 cells, operator 1 counted 810 cell and operator 2 counted 751 cells. **(B)** The proportion of cells with $n$ beads counted by the algorithm in the representative image (blue squares) or in whole slide photos averaged across 4 experiments (red triangles) compared to the model prediction (green circles). The latter two distributions are derived from Figure 3 in the main text.

than the number of cells counted by the human operators. This disparity arises because the algorithm counts multiple overlapping cells as one, as seen in the classification mask shown in Figure 1B.

Figure 2B shows the proportion of cells with $n$ beads automatically counted from the representative image (Figure 1) compared to the proportion of cells with $n$ beads automatically counted from whole slide photos averaged from 4 experiments (shown in Figure 3 in main text). There are disparities between these distributions. Thee disparities implies that to deduce properties of cell population dynamics, it is necessary to quantify every cell in the population and not sufficient to quantify a select few (even of 100s of cells).

The number of cells with $r$ red beads and $b$ blue beads (total $n = r + b$ beads) counted by the human operators is displayed in Figure 3. The algorithm does not distinguish between red and blue beads. From the proportion of cells with $n$ beads, say $p_n$ (shown in Figure 2), the expected proportion of cells with $r$ red and $b$ blue beads (total $n = r + b$), say $q_{r,b}$, qualitatively follows a binomial distribution:

$$\frac{q_{r,b}}{p_{r+b}} \approx \frac{1}{2^{r+b}} \binom{r+b}{r} = \frac{1}{2^{r+b}} \binom{r+b}{b} . \tag{1}$$

This is a prediction from our mathematical model.

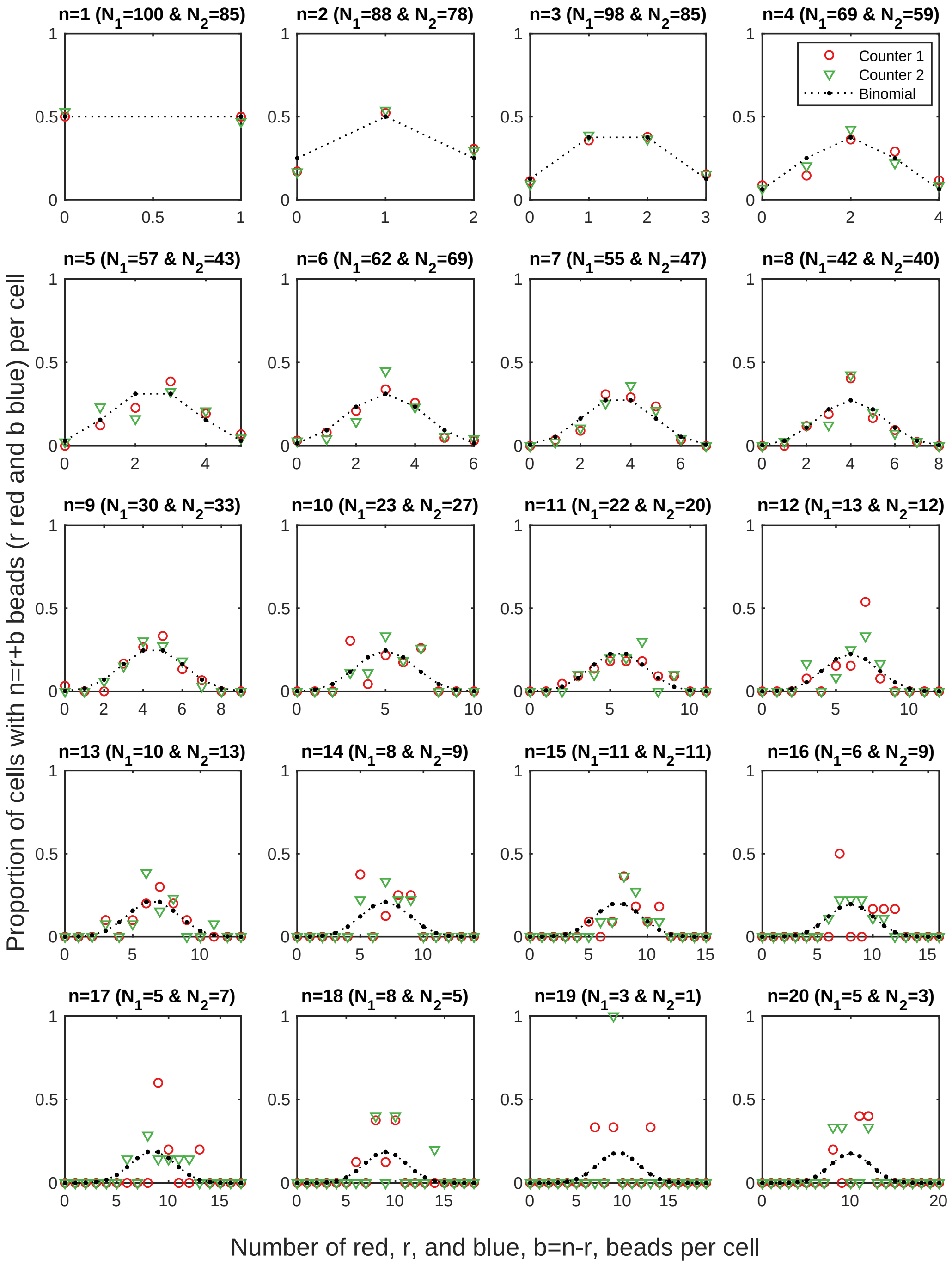


Figure 3: **Red and blue beads are binomially distirbuted within cells.** The proportion of cells with $n$ beads that contain $r$ red and $b$ blue beads ($n = r + b$) as counted by each human operator (red and green) compared to the binomial distribution (black) given by equation (1). Each plot displays the distribution of cells across $r$ (with $b = n - r$) for different $n$ values ranging from $n = 1$ to $n = 20$ beads (shown in the titles). The title of each plot also displays the number of cells with $n$ beads counted by each operator ($N_1$ and $N_2$).

### 2 Estimation of cell death rates

We can estimate the macrophage death rates from the total number of beads counted in each experiment displayed in Figures 3-5 in the main text. We assume that the death rate is constant throughout each

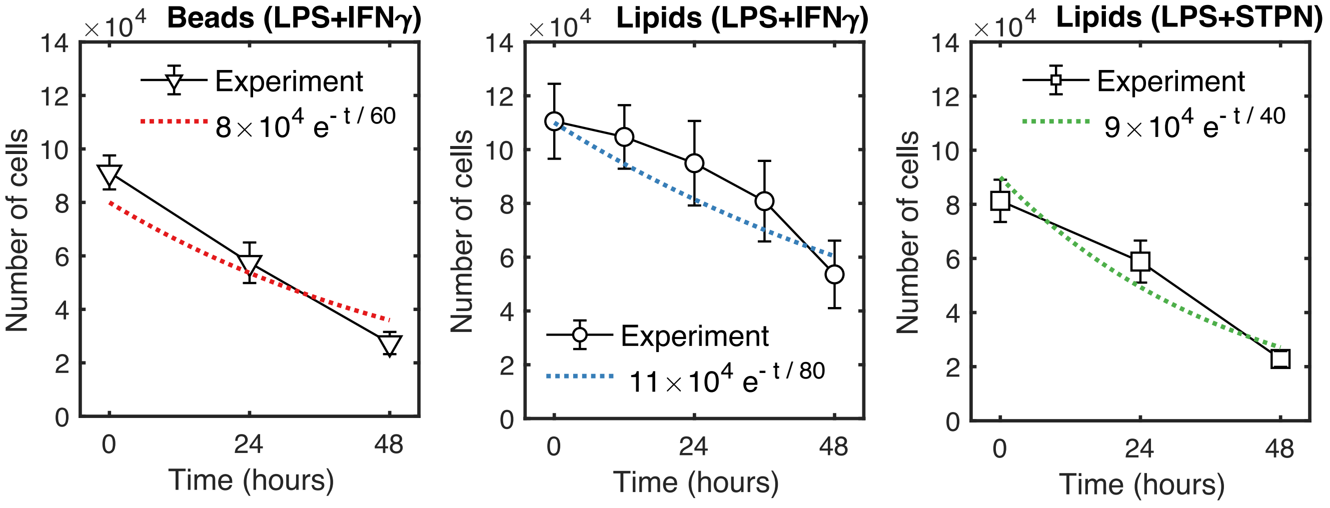


Figure 4: **Estimation of cell death rates.** Fit of an exponential function (equation (2)) to the change in cell numbers seen experimentally for bead loaded macrophages stimulated with LPS and IFN$\gamma$ (left), macrophages stimulated with LPS and IFN$\gamma$ (middle) and macrophages stimulated with LPS and STPN (right). The estimated parameters that represent the initial cell numbers $N_0$ cells and the cell death rate $\beta$ per hour are shown in the figure legends.

experiment. Under this assumption, the mathematical model predicts that the population size $N$ decays exponentially with time $t$ such that:

$$N(t) = N_0 e^{-\beta t} \ , \tag{2}$$

where $N_0$ cell is the initial number of cells and $\beta$ per hour is the mean cell death rate. Figure 4 shows a fit of this function to the experimental data with estimated parameter values of $N_0$ and $\beta$. For bead-loaded macrophages stimulated with LPS and IFN$\gamma$, we estimate that the death rate is $\beta = 1/60$ per hour. For macrophages stimulated with LPS and IFN$\gamma$, we estimate that the death rate is $\beta = 1/80$ per hour. For macrophages stimulated with LPS and STPN, we predict that the death rate is $\beta = 1/40$ per hour. These findings imply that bead loaded macrophages die 1.5 times as fast than those without beads and that LPS and STPN stimulated macrophages die twice as fast as those stimualted with LPS and IFN$\gamma$.

It is likely that the death rate does not remain constant throughout the experiment such that cell numbers do not experimentally decrease with time. The death rate is likely to increase with time as nutrients in the media are exhausted and apoptotic factors generated by macrophages, such as tumour necrosis factor, accumulates in the media.
